# Novel method for clarifying faster interstitial flow in the intimal side of the aorta under intraluminal pressurization

**DOI:** 10.1101/2021.07.09.451753

**Authors:** Wataru Fukui, Ujihara Yoshihiro, Masanori Nakamura, Shukei Sugita

## Abstract

Hypertension causes aortic thickening, especially on the intimal side. Although the production of the extracellular matrix is observed, the type of mechanical stress that produces this response remains unclear. In this study, we hypothesize that the interstitial flow causes the thickening. To validate this claim, we proposed a novel method to measure the velocity distribution in the radial direction in the aorta, which has been unclear. A fluorescent dye was introduced in the lumen of the mouse thoracic aorta *ex vivo*, intraluminal pressure was applied, and a time-lapse image in the radial-circumferential plane was acquired under a two-photon microscope. The flow of the fluorescent dye from the intimal to the adventitial sides in the aorta was successfully observed. The acquired image was converted to a radial-time image (i.e., kymograph), and the flow velocity was quantified by applying the one-dimensional advection-diffusion equation to the fluorescent images. The results revealed a higher interstitial flow velocity in the aortic walls under higher intraluminal pressure and a higher velocity on the more intimal side. Thus, the interstitial flow is a candidate for the mechanical stress causing hyperplasia of the aorta under hypertension.

## Introduction

Hypertension causes aortic thickening (Bevan, 1976), especially on the intimal side (Matsumoto & Hayashi, 1996). During aortic thickening, an increase in ground substances was observed (Matsumoto & Hayashi, 1996). Aortic thickening is caused by the response of the aorta to maintain the circumferential normal stress in the physiological state even under higher intraluminal pressures (Matsumoto & Hayashi, 1996). If the aortic walls are assumed to be incompressible, the intimal side is more stretched in the circumferential direction than the adventitial side under higher intraluminal pressure. Thus, the aortic response is considered to be triggered by this stretching. Specifically, smooth muscle cells (SMCs) are cyclically stretched in the circumferential direction with changes in the intraluminal pressure, and they are believed to sense the force and respond accordingly. Several *in vitro* studies have reported the upregulation of collagen synthesis: SMCs subjected to 20% strain produce more L-proline, which is essential for the synthesis of collagen, than those subjected to 10% strain (Reyna et al., 2004); stretched SMCs increase the expression of collagen α1 (Stanley et al., 2000); and SMCs under 18% strain show increased production of prolyl-4-hydroxylase subunit α-1, a catalyst for the hydroxylation of L-proline and collagen types I and III (Liu et al., 2015).

The interstitial flow in the aortic walls also increases under hypertension. The hydraulic conductance, calculated as the transmural interstitial flow velocity divided by the intraluminal pressure, was almost constant for intraluminal pressures of 75–150 mmHg (Baldwin & Wilson, 1993), indicating that a higher intraluminal pressure results in a faster interstitial flow in the aortic walls. When hydrostatic pressure is applied on the intraluminal surface of the aorta, liquid in the lumen enters the aortic walls (Vargas et al., 1979) and moves toward the adventitial side. Thus, SMCs in the aortic walls are subjected to fluid flow stress. Computer simulations have already indicated that SMCs are subjected to increased shear stress with increasing intraluminal pressure (Dabagh et al., 2008; Tada & Tarbell, 2002). *In vitro* studies have reported that SMCs respond to shear stress. For example, SMCs incubated on a culture dish and subjected to shear stress increase the production rate of nitric oxide (NO) and endothelial nitric oxide synthase (eNOS) (Kim et al., 2017), prostaglandin I_2_ and E_2_ (PGI_2_ and PGE_2_, respectively) (Wang & Tarbell, 2000), and platelet-derived growth factor (PDGF) and basic fibroblast growth factor (bFGF) (Sterpetti et al., 1994). SMCs embedded in collagen gels and subjected to shear stress also produce PGI_2_ and PGE_2_ (Wang & Tarbell, 2000). Interestingly, SMCs subjected to shear stress reduce the expression of contractile marker proteins such as smooth muscle myosin heavy chain, smoothelin, and calponin, indicating that an interstitial flow changes SMCs from a more contractile to a more synthetic state (Shi et al., 2010). Changes from the contractile to the synthetic phenotype of SMCs are associated with the increased synthesis of collagen and elastin (Sjölund et al., 1986). Therefore, the high shear stress due to the interstitial flow under hypertension might cause aortic thickening.

Previous studies measured the average interstitial flow in the aorta by measuring the liquid volume leaking out from the outer surface of the aorta and dividing it by the outer surface area (Baldwin & Wilson, 1993; Baldwin et al., 1997; Baldwin et al., 1992; Chooi et al., 2016; Deng et al., 1998; Gaballa et al., 1992; Lever et al., 1992; Lu et al., 2013; Nguyen et al., 2015; Shou et al., 2006; Tarbell et al., 1988; Wang et al., 2019). However, the interstitial flow velocity may differ at different locations in the aortic walls because these walls are heterogeneous. For example, collagen fibers are abundant on the dorsal and proximal sides relative to the ventral and distal sides (Concannon et al., 2020; Sugita & Matsumoto, 2013), and space in the dorsal side is smaller than that in the ventral side (Tedgui & Lever, 1987). Furthermore, smooth muscle-rich layers (SMLs) and elastic laminas (ELs), whose constituents are completely different, alternate in the radial direction.

If the interstitial flow influences aortic thickening, it must be higher in the intimal side. However, to the best of our knowledge, no method has been developed for measuring the local interstitial flow. Therefore, in this study, we proposed a novel method for quantifying the local interstitial flow velocity in the aorta. Then, we applied this method to measure the flow velocity distribution in the radial direction of the aorta.

## Materials & Methods

### Sample preparation

Five Slc:ddY mice (8–11 weeks, 30–42 g, Chubu Kagaku Shizai, Nagoya, Japan) were used as a test model. All animal experiments were approved by the Institutional Review Board of Animal Care at Nagoya Institute of Technology. The thoracic aorta was excised as reported in previous studies (Sugita et al., 2020; Sugita & Matsumoto, 2017). After the mouse was euthanized in a CO_2_ chamber, its thoracic aorta was exposed. As a length marker in the longitudinal direction *in vivo*, gentian violet dots were marked at 3-mm intervals on the surface of the aorta. After intercostal arteries were cauterized, the aorta was resected. The tubular sample was immersed in a buffer (0.5 mM CaCl_2_, 23.1 mM NaCl, 0.9 mM KCl, 0.2 mM MgSO_4_, 4.4 mM NaHCO_3_, 0.2 mM KH_2_PO_4_, 1.8 mM glucose) until the next experiment to maintain the activity of the SMCs in the aorta.

### Intraluminal pressurization

The obtained tubular sample was pressurized in a manner similar to that in previous studies (Sugita et al., 2020; Sugita & Matsumoto, 2017). Figure 1 shows a schematic of the experimental setup. The air pressure was regulated using an electropneumatic regulator (640BA20B, Asahi Enterprise, Tokyo, Japan) connected to a pressure source (0.3 MPa), a DA/AD converter (NI USB-6363, National Instruments, Austin, TX, USA), a personal computer (FMV BIBLO, Fujitsu, Tokyo, Japan; PC), and NI LabVIEW 2010 software (National Instruments). The pressure was applied to two reservoirs to convert air pressure to liquid pressure: one reservoir was filled with the buffer, and the other one was filled with 1.06 mM of uranine fluorescent dye solution (CI-45350, Tokyo Chemical Industry, Tokyo, Japan). An aortic sample was placed downstream of the reservoirs. Both ends of the aortic sample were tied to hypodermic needles (NN-2332R, Terumo, Tokyo, Japan) with suture threads (C-23-N2 7-0, Natsume Seisakusho, Tokyo, Japan), and the needles were fixed on a tissue bath. The sample was stretched in its longitudinal direction until the intervals of the length marker reached 3 mm. A pressure transducer (DX-300, Nihon Kohden, Tokyo, Japan) at the downstream side of the sample measured the pressure inside the tube through a strain amplifier (DPM-911B, Kyowa Electronic Instruments, Chofu, Japan), the DA/AD converter, and the software installed on the PC. The downstream side of the tube was connected to an outlet bath through a three-way valve (no. 3 in Fig. 1).

**Fig. 1.**
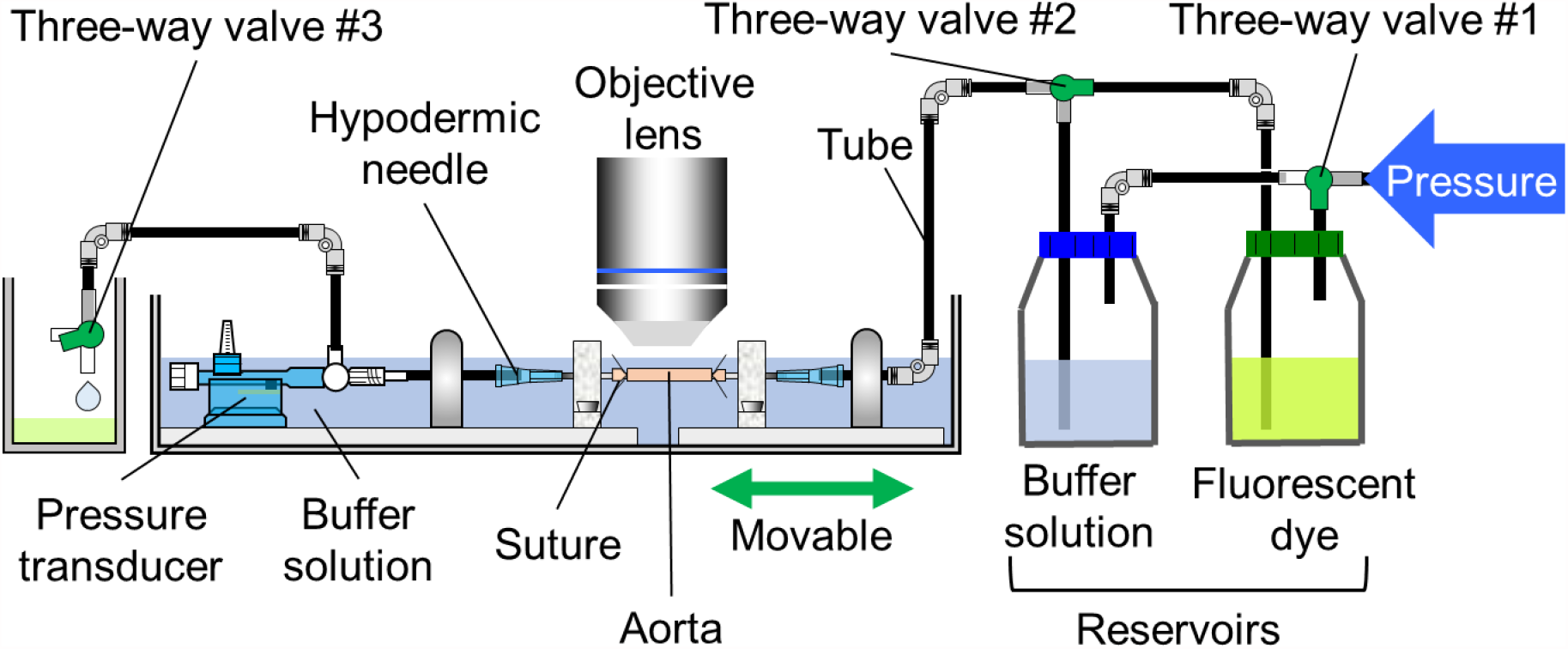
Schematic of experimental device for applying intraluminal pressure to a vessel. This device is a slightly modified version of the one fabricated in Sugita et al. (2017).

### Fluorescent microscopy

Fluorescent light was observed under a two-photon microscope (FV1200MPE, Olympus, Tokyo, Japan). A Ti:sapphire laser (wavelength: 800 nm, strength: 2.0%) was applied to the sample through a 60× objective lens (LUMPLFLN60XW, Olympus) and a V/G filter (FV-10-MRV/G, Olympus). To specify the position of the aorta, autofluorescence from the elastin in the aorta was mainly observed in a V-channel bandpass filter (420–460 nm). The fluorescent light of the fluorescent dye solution was imaged using a G-channel bandpass filter (495–540 nm).

### Experimental protocol

The intraluminal pressure *P* was set at 40 mmHg, and only the buffer was introduced into the aorta by switching three-way valves nos. 1 and 2 (Fig. 1). The flow rate was controlled such that the difference between the set and the measured pressures did not exceed 10 mmHg with three-way valve no. 3. The ELs were imaged in the plane perpendicular to the radial (*r*) direction, with an image size of 1 pixel (0.41 μm) in the longitudinal (*z*) and 512 pixels (211.97 μm) in the circumferential (*θ*) directions. By moving the objective lens in the *r*-direction in steps of 0.41 μm, the *r*-*θ* cross-sectional image was obtained.

The fluorescent dye solution was introduced into the aorta by switching three-way valves nos. 1 and 2. The aorta was imaged as stated above as a time-lapsed image with an interval of 2.7–3.5 s. Image capture was continued for 3–5 min. After imaging, the buffer solution was reintroduced into the aorta to wash out the fluorescent solution. The new position in the sample was then selected, and intraluminal pressurization and image capture were similarly repeated at 80, 120, and 160 mmHg.

All experiments were completed within 24 h after the resection of the sample.

### Image analysis

Image analysis was performed using image analysis software (ImageJ 1.52a, National Institutes of Health, Bethesda, MD, USA). Figure 2 shows a schematic of the image analysis process. The drift of the time-lapsed image stack obtained as described above was compensated by performing image correlation. The chord length and camber of the arch of any EL were measured, a circle was fitted to the EL, and its center coordinates were determined (Fig. 2a). By regarding the center coordinates as the origin of the polar coordinates, the image stack with polar coordinates was converted to an image stack with Cartesian coordinates (Fig. 2b). The converted radial direction still has an image resolution of 0.41 μm/pixel. The image stack was converted to time-lapsed images with 1-s intervals by linearly compensating every two sequential images.

**Fig. 2.**
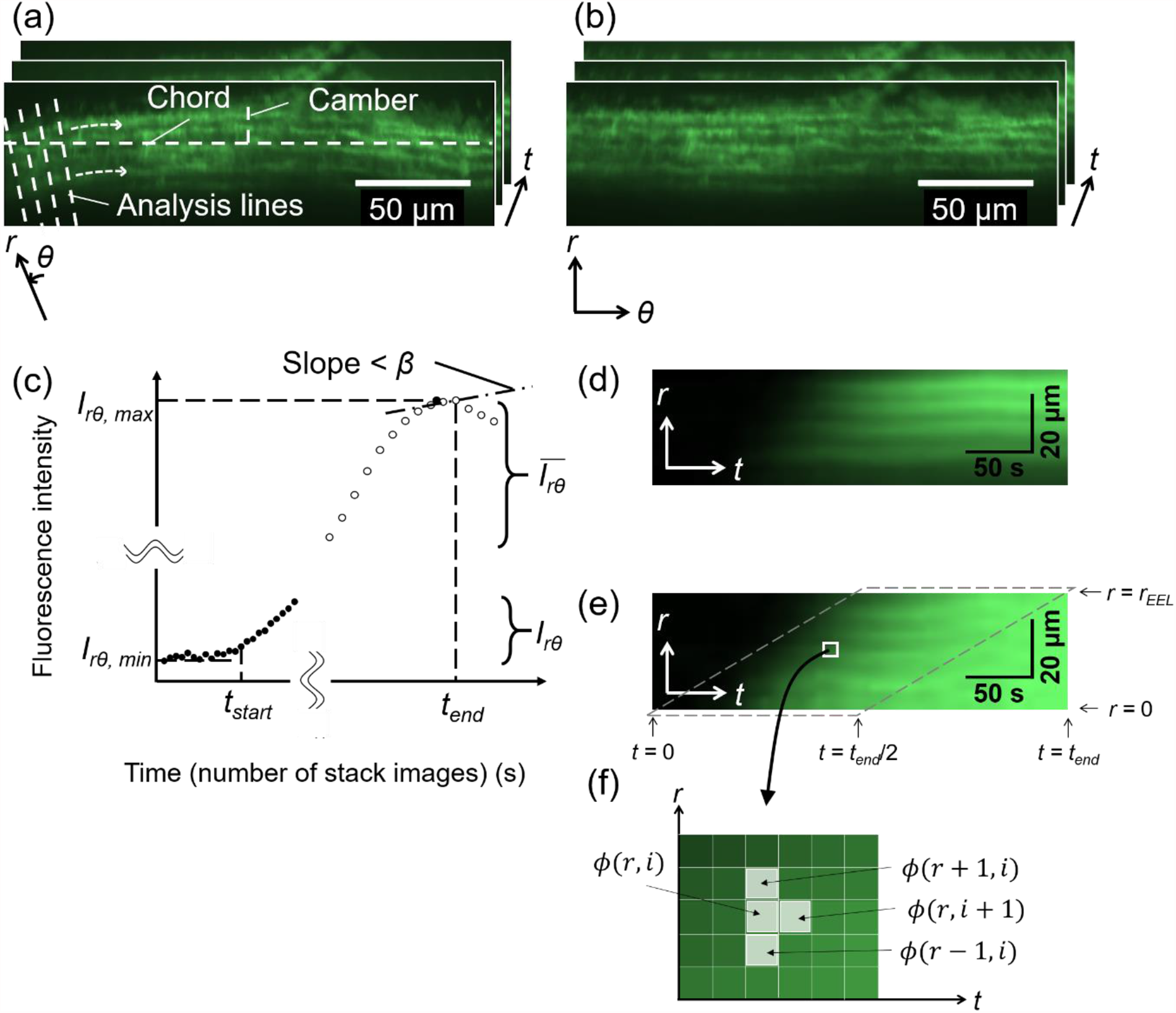
Schematic of image analysis. (a) An image stack of fluorescent dye captured in the radial-circumferential (*r*-*θ*) plane. (b) To convert the image of the aortic wall on an arch to images of a flattened one, the chord and camber lengths were measured from any arbitrary EL (outermost layer in this figure) and the center coordinates of the arc were determined. An analysis line was drawn from the center coordinates to an arbitrary position on the left edge of the images and rotated clockwise around the center coordinates. The fluorescence intensity on each line was recorded, and the image stack was reproduced from the intensity data. (c) Identification of start time *t*_*start*_ and end time *t*_*end*_. The average fluorescence intensity *I*_*rθ*_(*t*) in the aortic region of each slice was measured (closed circles), and the minimum *I*_*rθ,min*_ and maximum *I*_*rθ,max*_ were measured. *t*_*start*_ was defined as the time *t* when *I*_*rθ*_(*t*) slightly exceeded *I*_*rθ,min*_ (see main text for precise definition). The moving average of the fluorescence intensity 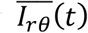 (open circles) was calculated, and *t*_*end*_ was defined as the time *t* when 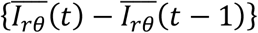 first became less than *β*. (d) A kymograph obtained by reslicing image (b) and averaging it on the *θ*-axis. (e) A normalized kymograph with 0 at *t*_*start*_ and 1 at *t*_*end*_. Normalization was performed at each *r*-coordinate. Further image analysis was performed in the region bordered by the broken line. (f) Magnified view of the kymograph. The differential of the intensity was calculated for each pixel. These images were taken at 80 mmHg.

There was a time delay until the fluorescent solution reached the sample position from the reservoir after the valves were opened. Thus, the start time *t*_*start*_ of the analysis was determined from the fluorescent intensity changes (Fig. 2c). First, the average intensity *I*_*rθ*_(*t*) of the image in the aortic region was calculated at each slice (*i*.*e*., each time *t*), and its maximum *I*_*rθ,max*_ and minimum *I*_*rθ,min*_ were determined. Then, the time *t* when {*I*_*rθ*_(*t*) − *I*_*rθ,min*_} first exceeded *α* times the difference between *I*_*rθ,max*_ and *I*_*rθ,min*_, that is,

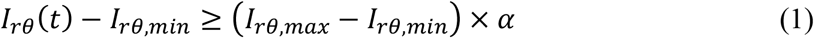

was determined as *t*_*start*_. Then, the moving average 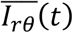 was calculated for time duration *t*_*MA*_ as

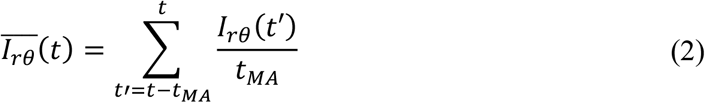

The end time *t*_*end*_ of the analysis was determined as the time when changes in 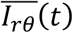 first became less than *β* after *t*_*start*_ (Fig. 2c). In this study, *α* = 0.5%, *t*_*MA*_ = 10 s, and *β* = 1 were selected. In the following analysis, only the time-lapsed images between *t*_*start*_ and *t*_*end*_ were used. The *r*-*θ* plane image stack was resliced into the *r*-*t* plane image stack to obtain the so-called kymograph stack, and this stack was then averaged in the *θ* direction (Fig. 2d). Images of only the media, from the internal to the external ELs, were cropped, and the image size was reduced to one-fifth the original for averaging the local intensities. The intensity *I*_*rt*_(*r, t*_*start*_) was subtracted from the intensity *I*_*rt*_(*r, t*) at the coordinates (*r, t*) in the kymograph to remove the background value as follows:

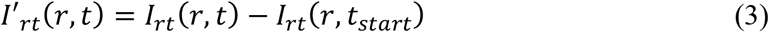

Then, the obtained intensity *I*′_*rt*_(*r, t*) was normalized by the intensity *I*′_*rt*_(*r, t*_*end*_) at *t*_*end*_ as follows (see Fig. 2e):

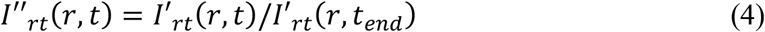

This process was performed to eliminate the effect of the difference in the intensity in the radial direction because the light at deeper tissues (intimal side) tends to be weak and dispersed owing to the long distance from the objective lens. The obtained intensity *I*′*′*_*rt*_(*r, t*) in the *r*-*t* plane was used in the following analysis.

### Calculation of velocity and diffusion coefficient

To measure the interstitial flow in the *r*-axis, the following one-dimensional advection-diffusion equation was used:

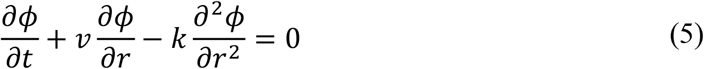

Here, *ϕ* is the concentration of the fluorescent solution; *v*, the interstitial flow velocity; and *k*, the diffusion coefficient. The discretization of Eq. (5) gives

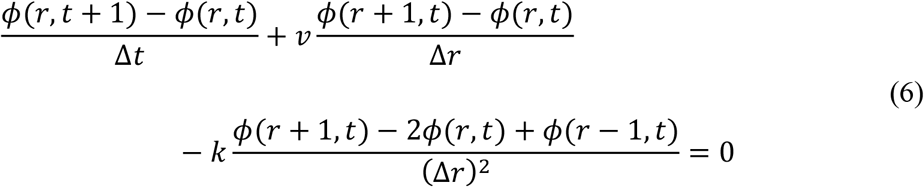

where *ϕ*(*r, t*) is the concentration at a coordinate (*r, t*) in the kymograph (see Fig. 2f). The coefficient of each term in Eq. (6) is expressed as

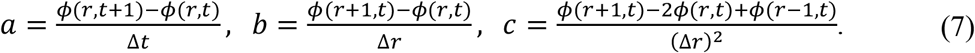

Eq. (6) is then expressed as

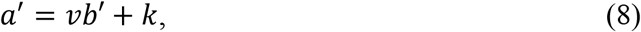

where

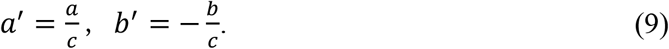

The intensity in the normalized kymograph shown in Fig. 2e was considered to reflect the concentration of the fluorescent dye (see Supplementary Material S1):

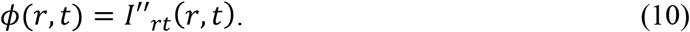

Further, the application of Eq. (6) to each pixel in the kymograph produced many combinations of (*a’, b’*). These combinations were plotted on a graph with ordinate *a’* and abscissa *b’*. To eliminate the small intensity changes that often produce extremely high values in the calculation of the differentiation, the kymograph in the ranges with

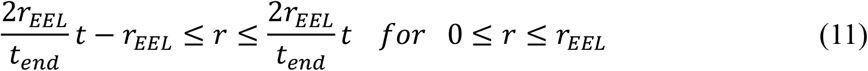

were used for further analysis, where *r*_*EEL*_ is the radial coordinate at the external elastic lamina (EEL, Fig. 2e). The extreme data (*a’, b’*) were excluded using the modified Stahel-Donoho method (Wada, 2010) in R software (v. 3.5.1, University of Auckland, Auckland, North Island, NZ). A linear line was fitted to the plots (*a’, b’*) with a least-squares regression, and the slope and *b’*-axis intercept were determined as *v* and *k*, respectively, by using Eq. (8).

### Distribution of interstitial flow velocity in radial direction

The distribution of the interstitial flow velocity in the radial direction was measured. In the kymograph image, a local area with a length of 5–10 pixels (ca. 2–4 µm) in the radial direction was selected in the EL and SML regions for the autofluorescence image of elastin. From the intimal side, these ELs were named EL1, EL2,, and EL3, and the SMLs were named SML1, SML2, and SML3. Then, the abovementioned process was applied to the image to obtain *v*, except that the image size was reduced to one-fifth the original for averaging the local intensities.

To determine the radial position of the local area image, the normalized distance *d* from the internal elastic lamina (IEL) was calculated as the distance from the IEL divided by the distance between the IEL and the EEL.

### Statistical method

The correlation coefficients *R* between plots such as *a’*-*b’, v*-*P, k*-*P*, and *v*-*d* were tested using Student’s *t*-test. The velocity difference between SMLs was tested using the Steel-Dwass test. Data were expressed as mean ± SD. A significance level of *p* = 0.05 was used.

## Results

### Observation of fluorescent solutions during intraluminal pressurization

Figure 3 and Movie 1 show raw time-lapsed images of the fluorescent solution in the thoracic aorta in the *r*-*θ* plane under intraluminal pressurization. After the fluorescent solution reached the aorta from the reservoir, a high-intensity area spread from the bottom (i.e., intimal side) to the top (i.e., adventitial side). The high intensity reached the adventitial side more quickly under a higher intraluminal pressure. Although this is the fluorescent dye channel image and not the main elastin autofluorescence channel, the EL was clearly observed after the fluorescent dye reached the aortic walls.

**Fig. 3.**
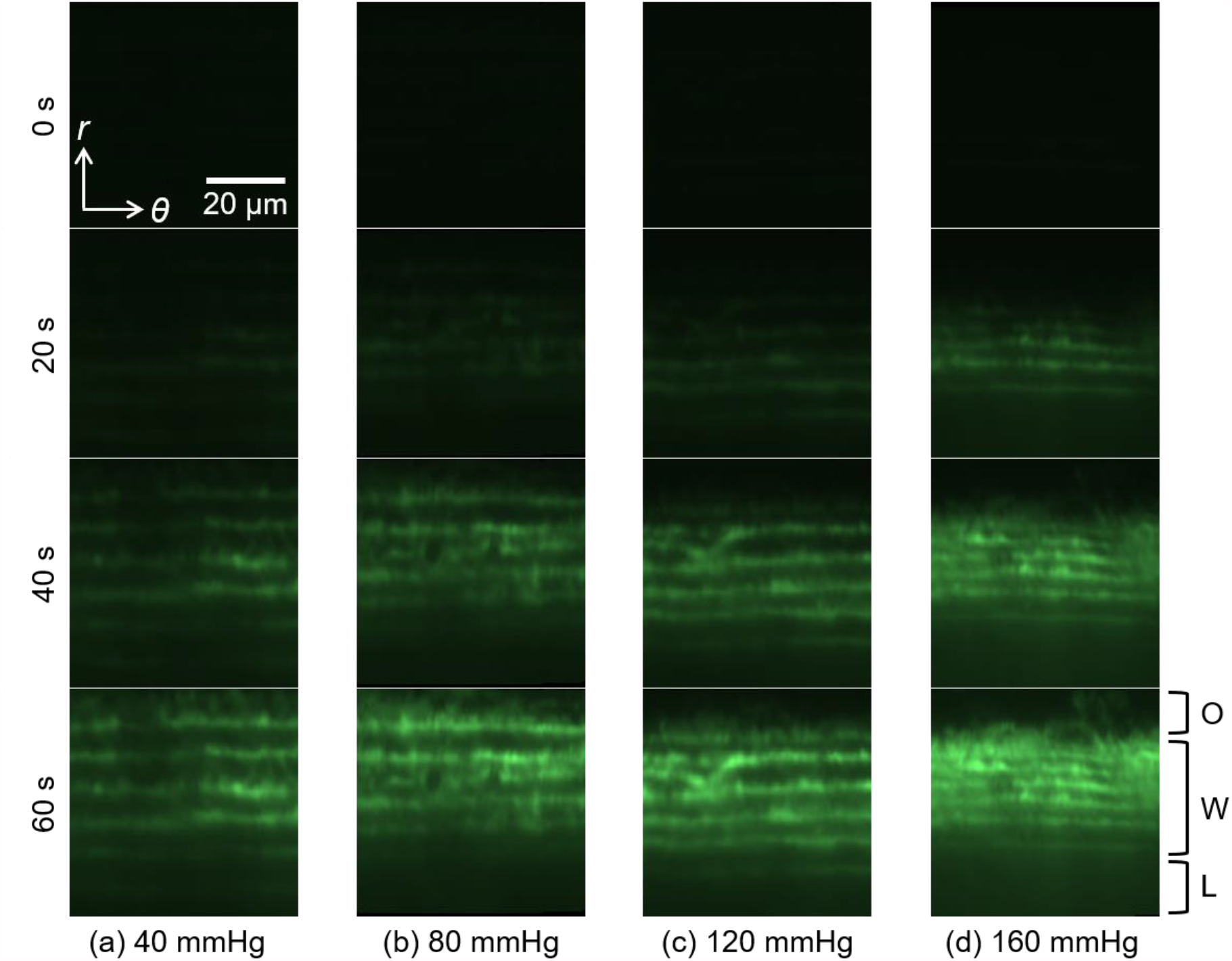
Time-lapsed images of fluorescence in the aortic walls in the radial-circumferential (*r*-*θ*) cross-section at intraluminal pressures of (a) 40, (b) 80, (c) 120, and (d) 160 mmHg at 0, 20, 40, and 60 s after *t*_start_. From the top to the bottom sides, the outside of the aorta (O), aortic wall (W), and lumen (L) are shown.

Figure 4 shows the kymographs at 40, 80, 120, and 160 mmHg. These kymographs show the movement of the higher intensity from the intimal to the adventitial side, indicating that the fluorescent dye flows through the aortic walls. Furthermore, the time length of the kymograph tended to be shorter at higher intraluminal pressures, indicating that the fluorescent intensity at higher pressure reached a stable state more quickly. In this kymograph, stripe patterns were also seen under all pressure conditions. The higher-intensity regions in the strip patterns correspond to the region of ELs.

**Fig. 4.**
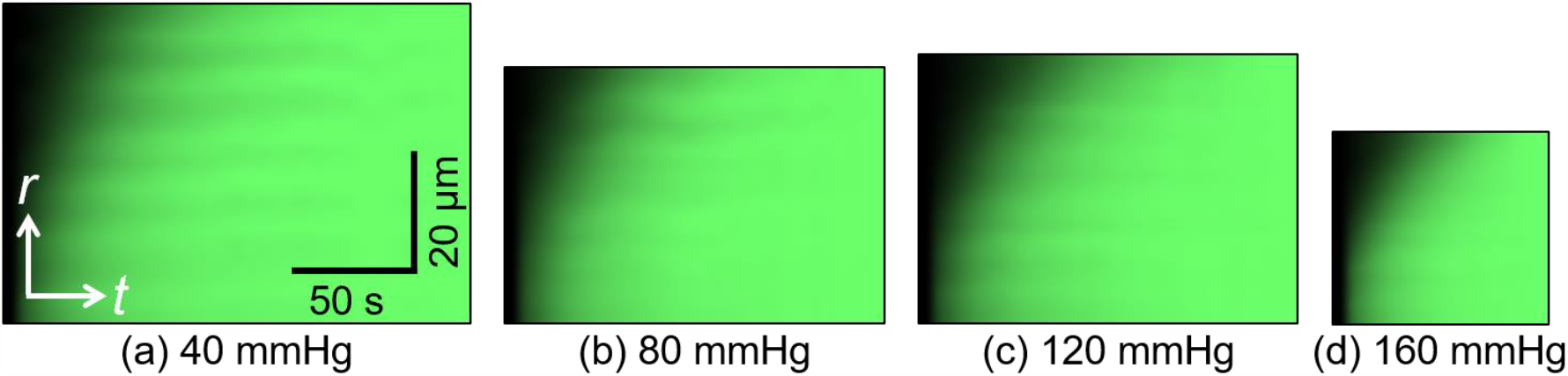
Kymograph of fluorescent dye solution at intraluminal pressures of (a) 40, (b) 80, (c) 120, and (d) 160 mmHg. *t*, time axis; *r*, radial axis. These images were obtained from the images shown in Fig. 3.

### Interstitial flow velocity and diffusion coefficient in analysis of whole thickness

Figure 5 shows typical graphs of combinations of (*a’, b’*). The density of the plots was high from the bottom left to the top right, indicating that the velocity expressed as the slope of the fitted line is positive. The slope of the fitting line increased with increasing pressure, as given by *a’* = 0.06*b’* + 0.12 at 40 mmHg (Fig. 5a), *a’* = 0.28*b’* + 0.19 at 80 mmHg (Fig. 5b), *a’* = 0.40*b’* + 0.34 at 120 mmHg (Fig. 5c), and *a’* = 0.59*b’* + 0.98 at 160 mmHg (Fig. 5d). The number of plots tended to decrease at higher intraluminal pressures because the image sizes of the kymograph under such conditions were small (Fig. 4).

**Fig. 5.**
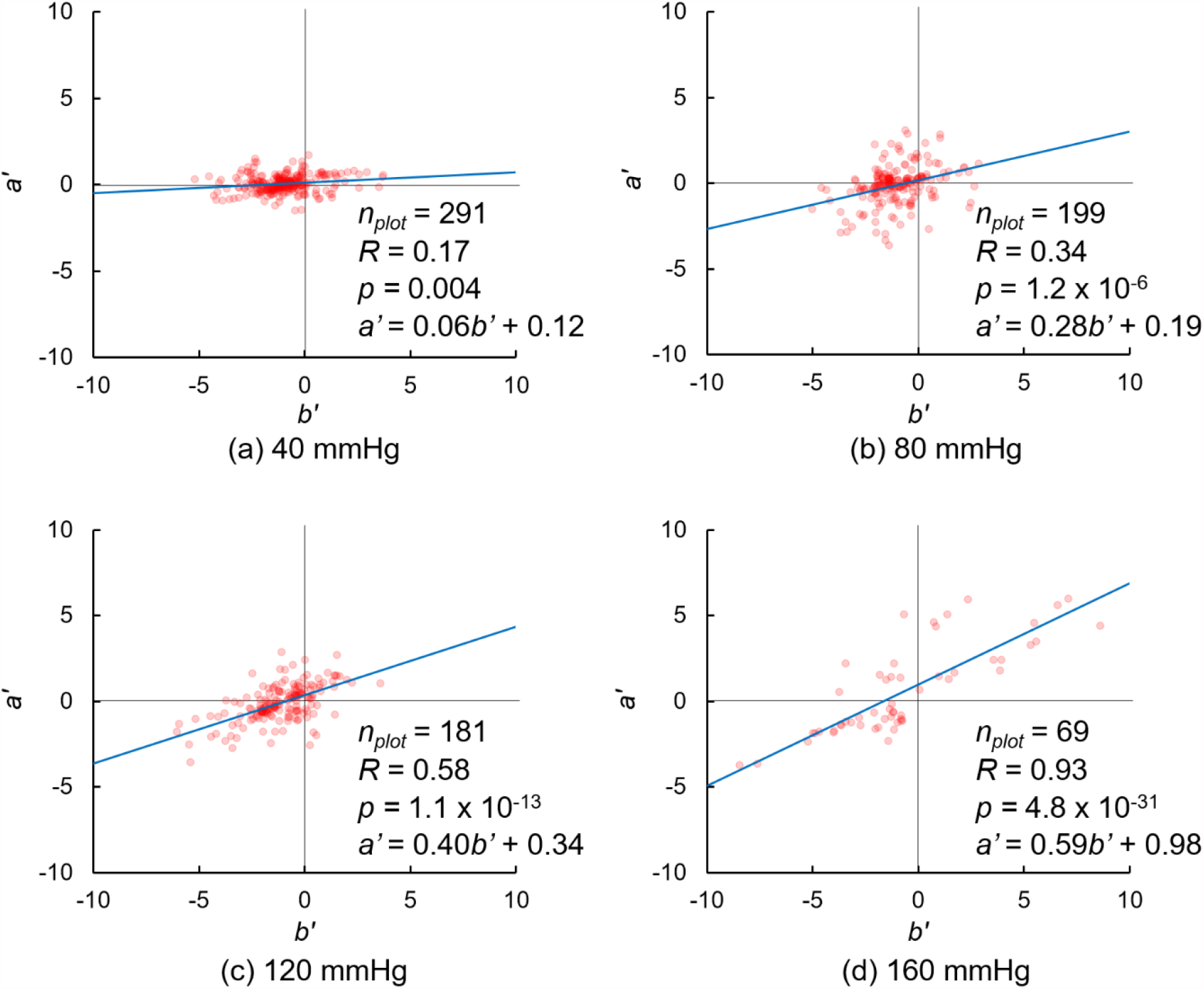
Plots of *a’* and *b’* as given by Eq. (8) at intraluminal pressures of (a) 40 mmHg, (b) 80 mmHg, (c) 120 mmHg, and (d) 160 mmHg. *n*_*plot*_, number of plots. These graphs are obtained from the images shown in Figs. 3 and 4.

Figure 6 shows plots of the interstitial flow velocity *v* and diffusion coefficient *k* against the pressure *P*. Because a significant correlation between *a’* and *b’* was not obtained from data at 80 mmHg and the adventitial side was destroyed owing to the laser as indicated by the data at 160 mmHg, those data were not used in the following analysis. The velocities *v* were 0.23 ± 0.16 μm/s at 40 mmHg (*n* = 13), 0.34 ± 0.15 μm/s at 80 mmHg (*n* = 10), 0.40 ± 0.23 μm/s at 120 mmHg (*n* = 11), and 0.40 ± 0.27 μm/s at 160 mmHg (*n* = 10). Further, the diffusion coefficients *k* were 0.24 ± 0.18 μm^2^/s at 40 mmHg (*n* = 13), 0.41 ± 0.22 μm^2^/s at 80 mmHg (*n* = 10), 0.35 ± 0.29 μm^2^/s at 120 mmHg (*n* = 11), and 0.56 ± 0.47 μm^2^/s at 160 mmHg (*n* = 10). Both *v* and *k* increased with increasing pressure *P*, and the correlation coefficients for the *P*-*v* and *P*-*k* relationships were significant.

**Fig. 6.**
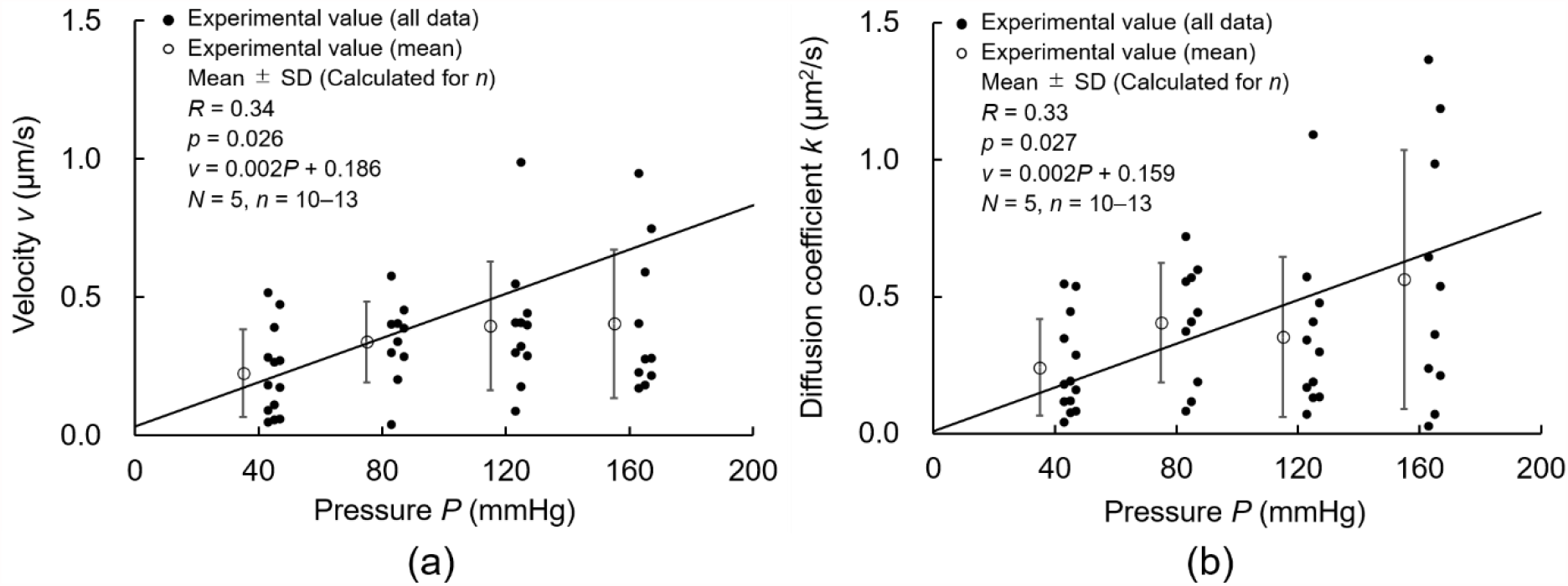
Correlations between intraluminal pressure *P* and both (a) interstitial flow velocity *v* and (b) diffusion coefficient *k. N*, Number of mice; *n*, Number of data; *R*, correlation coefficient.

### Velocity distribution in radial direction

Figure S2.1 shows the kymograph in the local EL and SML regions. The intensity change in the radial direction was negligible, whereas that in the time direction was recognizable. Figure S2.2 shows the plots of *a’* and *b’* in the EL and SML regions. Significant correlations were obtained, as seen in the analysis for the whole wall thickness (Fig. 5). The slope of the relationship between *a’* and *b’* was steeper in the SML region and intimal side. Figure S2.3 shows the slope, that is, the interstitial flow velocity *v*. In ELs, these velocities were lower or sometimes negative. Thus, the reliability of the velocity in ELs was not enough, and only the SML data were used. Figure 7 shows the interstitial flow velocity *v* in the SMLs plotted to the normalized radial position *d* for all pressure levels. Negative and significant correlations were found at 80–160 mmHg, whereas the *p*-value of the relationship at 40 mmHg was slightly larger than 0.05. Figure 8 shows the interstitial flow velocity *v* in SML1–SML3. At 120 and 160 mmHg, *v* in SML1 was significantly higher than that in SML2 and SML3. At 40 and 80 mmHg, *v* in SML1 tended to be higher than that in other SMLs, and significant differences were not seen. These results indicate that the interstitial flow velocity is higher on the intimal side than on the adventitial side.

**Fig. 7.**
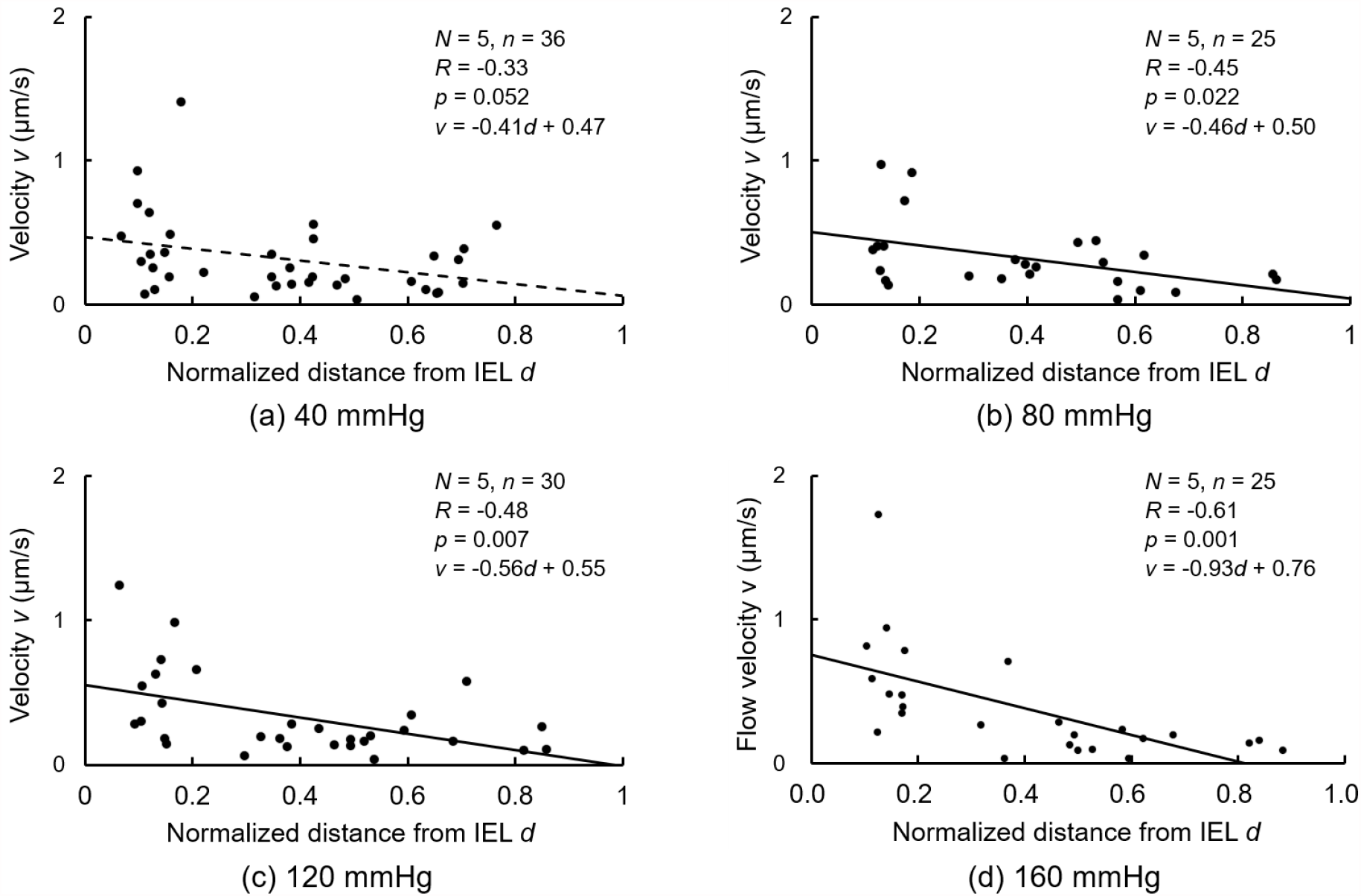
Correlations between normalized distance *d* from the IEL and interstitial flow velocity *v* at intraluminal pressure *P* of (a) 40, (b) 80, (c) 120, and (d) 160 mmHg. *N*, Number of mice; *n*, Number of data; *R*, correlation coefficient.

**Fig. 8.**
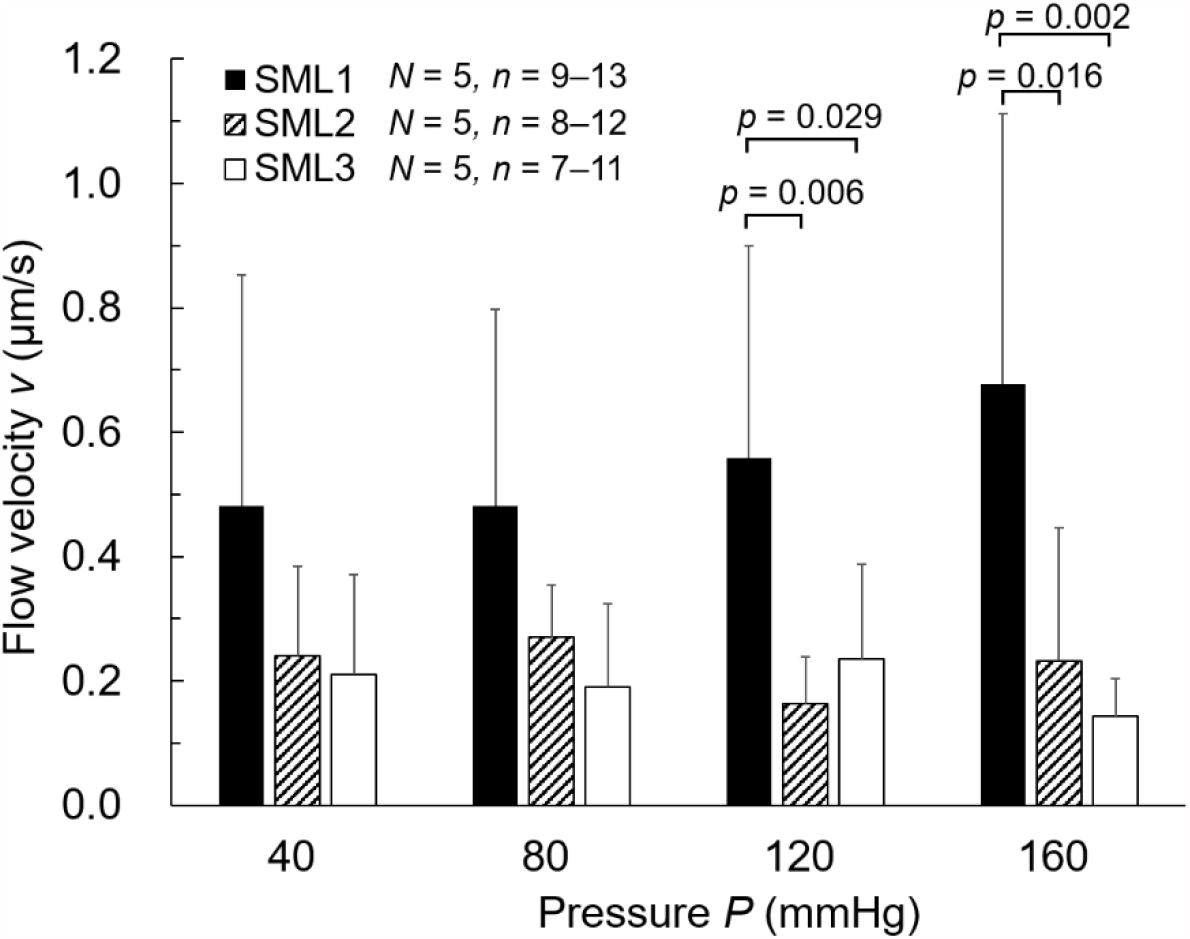
Comparison of flow velocity *v* between SML1–SML3. *N*, Number of mice; *n*, Number of data.

## Discussion

In this study, we proposed a novel method to quantify the local interstitial flow velocity in the aortic wall. Upon applying an intraluminal pressure to the mouse thoracic aorta, the fluorescent dye solution introduced in the lumen was observed to move from the intimal to the adventitial side under a two-photon microscope. By applying a one-dimensional advection-diffusion equation to time-lapsed images of the fluorescence intensity, the flow velocity and diffusion coefficient were successfully obtained. This study makes the following main contributions: (1) it proposes a novel method to observe the flow through the change in the concentration of the fluorescent dye, (2) it directly observes the higher interstitial flow velocity in the radial direction under higher intraluminal pressure, and (3) it reveals the higher interstitial flow velocity in the intimal side of the aorta.

The interstitial flow in the aorta increased with increasing pressure at its lumen, in keeping with previous studies (Baldwin & Wilson, 1993; Baldwin et al., 1992). In these studies, the hydraulic conductance was almost constant regardless of the intraluminal pressure, indicating that the flow velocity increased with an increase in the intraluminal pressure. The interstitial flow was affected by not only the pressure difference but also the changes in the aortic thickness. A decrease in the wall thickness, which was measured as the distance between the IEL and the EEL, was also found with increased intraluminal pressure (Fig. S3.1). Thus, we concluded that the increase in the pressure and the decrease in the wall thickness both increase the interstitial flow velocity.

This study directly and experimentally demonstrated higher interstitial flow velocity in the intimal side of the SML (Fig. 7). When the continuum theory is applied to the inner and outer surfaces, the interstitial flow velocity at the inner surface must be higher than that at the outer surface, because the outer surface area is larger owing to the cylindrical shape of the aorta. However, the maximum rate of decrease in the velocity was estimated to be only 8%. The curvature radius of the ELs at the intimal and adventitial sides was ~990 µm and ~1080 µm, respectively; in other words, the intimal surface area was 8% smaller than the adventitial surface area at 40 mmHg. The fitting lines in Fig. 7 indicate a much larger rate of decrease at the outer surface compared to that at the inner surface. Thus, other factors must influence the higher interstitial flow velocity in the more intimal side of the aorta. Through simulations, Huang et al. (1994) revealed much faster flow at the fenestral pores in the IEL, and Tada and Tarbell (2000, 2002) predicted that the flow throw the pores was jet flow. Thus, fenestral pores might influence the faster flow in the intimal side.

Hypertensive rats showed aortic thickening, especially in the intimal side (Matsumoto & Hayashi, 1996); this might have been caused by the faster interstitial flow in the intimal side, as shown in this study. SMCs in collagen gel subjected to a pressure of 0.005 Pa reduced the production of contractile-type proteins, indicating that they changed into a synthetic phenotype (Shi et al., 2010). Changes to the synthetic type improve the synthesis of collagen (Sjölund et al., 1986). Because an increase in ground substances was also seen in hypertensive rats (Matsumoto & Hayashi, 1996), the higher shear stress induced by faster fluid flow might be a cause of the aortic thickening seen in hypertension. This issue requires further investigation.

In this study, the interstitial flow velocity in the *ex vivo* mice aorta was respectively measured to be 0.34 and 0.40 µm/s at 80 and 120 mmHg on average; these values were higher than previously reported ones. The velocity in *ex vivo* aortas at 100 mmHg was reported as 0.055 µm/s (Vargas et al., 1979), 0.052 µm/s, (Baldwin et al., 1992), and 0.045 µm/s (Baldwin & Wilson, 1993) in rabbits and ~0.03 µm/s (Shou et al., 2006), 0.025 µm/s (Nguyen et al., 2015), and ~0.03 µm/s (Toussaint et al., 2017) in rats. Various factors might cause these velocity differences. The most likely factor is that in this study, unlike in previous studies, the adventitia was removed to observe the fluorescence in the media. Because the adventitia seems to resist the interstitial flow, their removal might result in the higher interstitial flow in this study. The second-most likely factor is the absence of endothelial cells (ECs). The interstitial flow velocity with ECs is higher than that without ECs (Baldwin & Wilson, 1993; Lu et al., 2013; Shou et al., 2006; Tedgui & Lever, 1984; Toussaint et al., 2017). Baldwin and Wilson (1993) reported that the interstitial flow velocity with ECs was 2.2 times larger than that without ECs at 100 mmHg. Although we have not confirmed the existence of ECs in the present study, ECs might have been removed by the experimental process. For example, the introduction of air into the aorta (Ralevic et al., 1989) and the buckling of the IEL owing to the removal of the intraluminal pressure (Baldwin et al., 1982) can detach the ECs. In the present study, the intraluminal side of the aortic specimen was exposed to the air and depressurized when the aorta was isolated from the animal body. These processes might peel the ECs off, thereby increasing the velocity. The third-most likely factor is the viscosity of the solution. Tarbell et al. (1988) reported that the viscosity of the Krebs solution is 0.81 times that of a solution containing 4% bovine serum albumin, and the interstitial flow without albumin is 1.8 times faster at 110 mmHg. Thus, the interstitial flow without albumin might have increased velocity in this study. Considering the previous two factors, namely, the effects of EC removal and the existence of albumin, the magnitude of the interstitial flow velocity was considered comparable to that of previously reported velocities. The other possible factors, though these are considered unlikely, are the differences in the animals and the laser scanning. With regard to the effect of the laser, scanning using the two-photon microscope was performed from the intimal to the adventitial side at a speed of 25 µm/s in the radial direction. The interstitial flow velocity for all data in this study was 0.39 µm/s; this results in a 1.6% error in the interstitial flow velocity.

Although the abovementioned factors might have resulted in the higher interstitial velocity in our study, the actual flow velocity in the aorta might be higher than the interstitial velocity in previous studies. Previous studies reported the averaged interstitial velocities determined by dividing the flow rate by the area of the intraluminal surface. However, the actual area where liquid can flow in the aorta is smaller owing to the existence of the extracellular matrix and cells. Thus, the interstitial flow velocities measured in the previous studies underestimate the actual liquid flow velocity. Tedgui and Lever (1987) reported that the extracellular space was ~40% in the media at 70 mmHg. If the area in which the liquid can flow in the radial direction is 40% of the total area of the intraluminal surface, the actual flow velocity is 2.5 times the interstitial velocity measured in previous studies. However, differences still exist between the interstitial flow in previous studies and our study. Therefore, this issue will also require further investigation.

The proposed method to measure the interstitial flow velocity differs from previous methods, in which the total volume of the liquid eluted from the aorta was measured and divided by the surface area, in several ways (Baldwin & Wilson, 1993; Baldwin et al., 1997; Baldwin et al., 1992; Chooi et al., 2016; Deng et al., 1998; Lever et al., 1992; Lu et al., 2013; Nguyen et al., 2015; Shou et al., 2006; Tarbell et al., 1988; Wang et al., 2019). First, it can directly observe the interstitial flow in the aorta. Thus, the local difference in the interstitial flow can be investigated in the *z*-, *θ*-, and even *r*-directions, unlike in previous studies. Specifically, because the structure of ELs and SMLs is quite different, there might be a big difference between SMLs and ELs that might affect the local velocity in the *r*-direction. Second, it can measure the velocity for a limited time; by contrast, previous methods do not have a time limitation. Because the measurements using the proposed method depend on changes in the concentration of the fluorescent dye solutions, they cannot be performed after the concentration reaches a stable value.

The interstitial flow velocity *v* was obtained under several assumptions. First, we assumed a constant flow velocity during the experiment. Further, Lever et al. (1992) reported that the velocity decreased with time for ~10 min after intraluminal pressurization. Because the present study measures the velocity for a few minutes after pressurization, the interstitial flow velocity in the stable state might be lower than that observed in this study. The aorta is viscoelastic, and we have often observed its outward movement during pressurization. However, the aortic thickness remains almost constant (Fig. S3.1), and the effect of the viscoelasticity of the aorta should be negligibly small. Second, we considered that the fluorescent intensity reflected the concentration of the fluorescent dye solution. This is because the investigation of the relationship between the intensity and the concentration showed a significant and strong correlation (see Supplementary Materials S1), indicating that the intensity can be used as a concentration index.

An EL-like structure, namely, a stripe pattern, was observed in the fluorescent dye solution image (Fig. 3). Because a high intensity was not observed at the start time *t*_*start*_, this should not be the autofluorescence from elastin. When an aorta was sectioned perpendicular to the longitudinal direction and immersed in the fluorescent dye solution without pressurization, following which the dye was washed out with the buffer, a higher intensity was observed in ELs than in SMLs (see Supplementary Materials S4). Therefore, we assume that more fluorescent dye attaches to something in ELs rather than that in SMLs.

The proposed method can measure the diffusion coefficient *k* of the fluorescent dye in the aortic walls (Fig. 6b). Notably, even though both flow and diffusion conveyed the fluorescent dye from the lumen to the adventitial side, the flow, and not the diffusion, mainly conveyed fluorescent dyes because the Peclet number was much larger than 1 (see Supplementary Materials S5). Although the movement of the fluorescent dye by diffusion was much smaller than that by flow, the measured diffusion coefficient *k* seemed appropriate because the diffusion coefficient *k* under the condition without a flow showed a similar value to that under the condition with a flow (see Supplementary Materials S6). Interestingly, the diffusion coefficient increased with an increase in the intraluminal pressure. Tedgui and Lever (1987) reported that the porosity of the liquid to fill the media at 70 mmHg was lower than that at 180 mmHg. This indicates that voids in the media increased with increasing intraluminal pressure, resulting in a higher diffusion coefficient as obtained in this study.

## Conclusion

We proposed a novel method for directly observing the flow of a fluorescent dye and for quantifying the local interstitial flow velocity in the aortic wall. As a result, a higher interstitial flow in the radial direction was confirmed under the higher intraluminal pressurization of the aorta. Moreover, the interstitial flow was faster at the more intimal side of the aorta.

## Acknowledgements

This work was supported in part by AMED Grant Number JP20gm0810005, JSPS KAKENHI Grant Number 21H04955a, and Tatematsu Foundation. We would like to thank Editage (www.editage.com) for English language editing.

## Supplementary Materials

### S1. Can the fluorescent intensity be used as an index of the concentration of fluorescent dye?

This study experimentally investigated whether the fluorescent intensity could be used as an index of the concentration of the fluorescent dye.

We prepared a fluorescent dye solution (CI-45350, Tokyo Chemical Industry, Tokyo, Japan) with a concentration of 1.06 mM, corresponding to the solution used in this study, as well as lower concentrations of 0.53, 0.33, 0.27, 0.21, 0.18, 0.15, and 0.13 mM. Figure S1.1 shows a schematic of the experimental setup. First, 50 µL of the solution was dropped into a large Petri dish (MS-11600, Sumitomo Bakelite, Tokyo, Japan), and then, a smaller Petri dish (MS-11350, Sumitomo Bakelite, Tokyo, Japan) was placed on the solution. These dishes were placed on the stage of a two-photon microscope (FV1200MPE, Olympus, Tokyo, Japan), and the fluorescent dye was observed using the G-channel bandpass filter (495–540 nm). An image stack with a size of 512 × 512 pixels (212 × 212 µm) in the *x*-*y* plane and 30 µm with every 5 µm in the *z*-direction (optical direction) was captured. The captured image was resliced from the *y*-direction, and a 2D *x*-*z* projection image was obtained as the maximum intensity. Then, the image in the *x*-direction was averaged, and the maximum value in the *z*-direction was defined as the intensity of the fluorescent solution.

Figure S1.2 shows the relationship between the fluorescent intensity and the concentration. The intensity is significantly and highly correlated with the concentration (*R* = 0.999, *p* < 0.01). This result indicates that the concentration can be evaluated from the intensity of the fluorescent image.

**Fig. S1.1.**
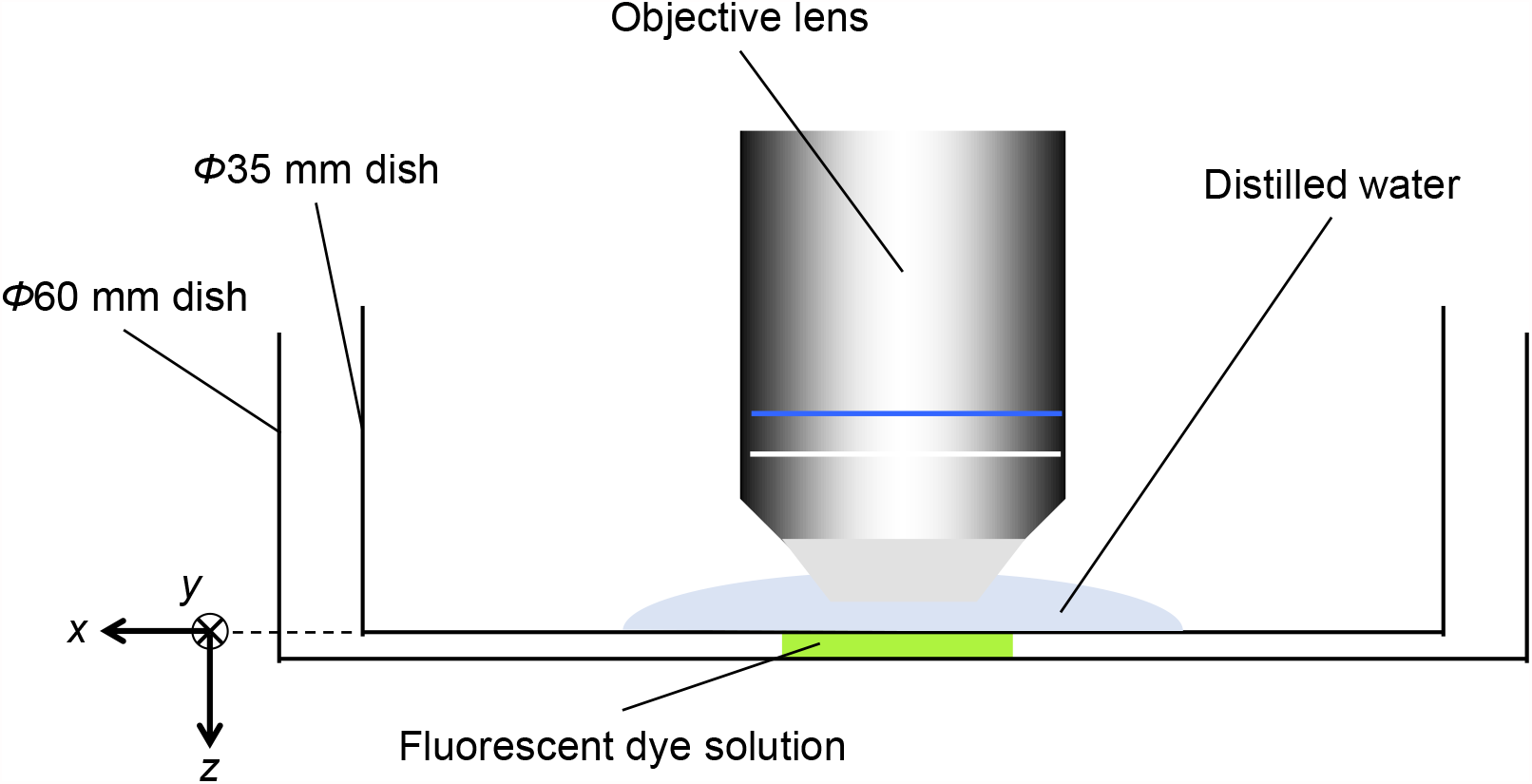
A schematic illustration of intensity measurement of fluorescent dye.

**Fig. S1.2.**
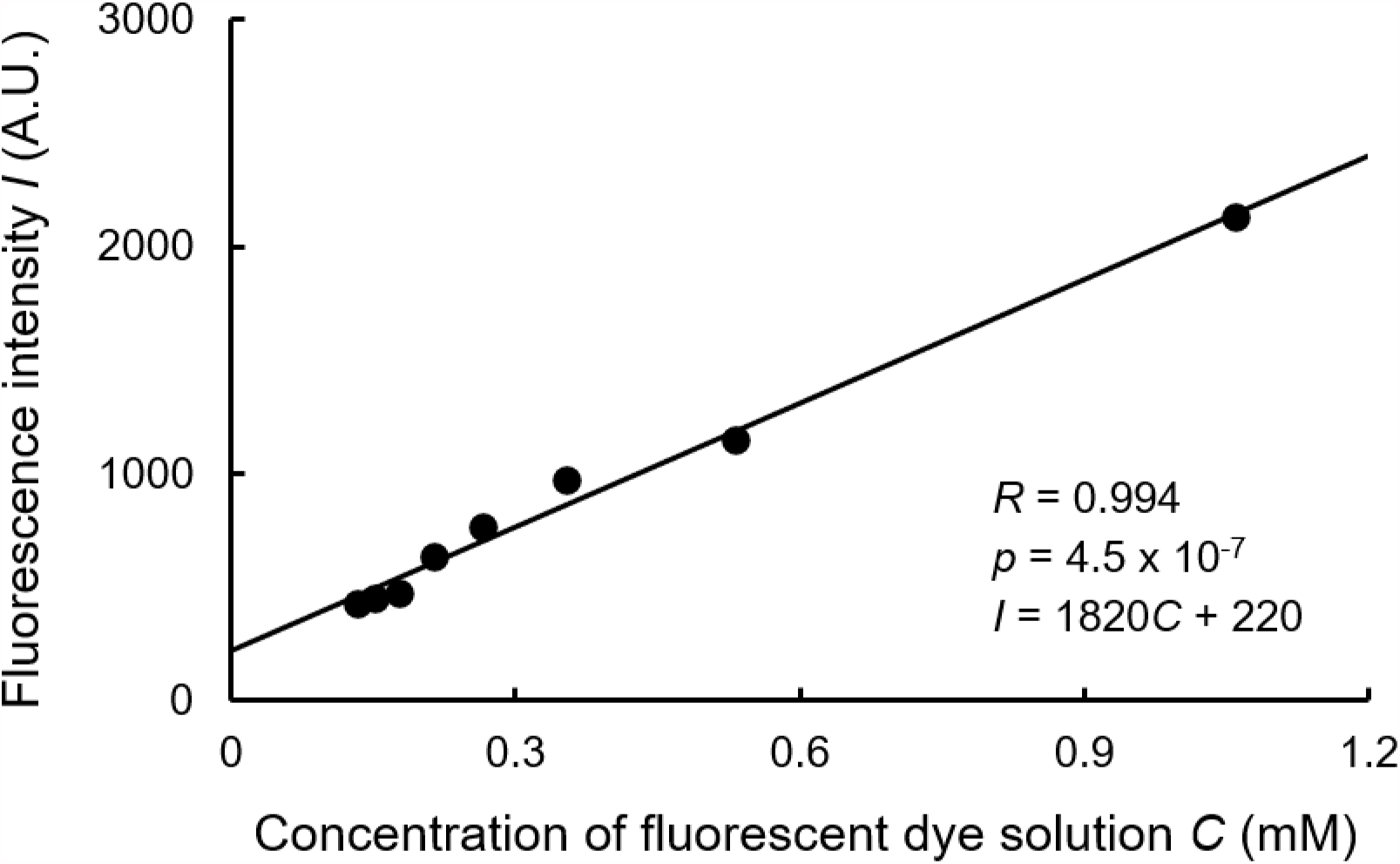
Fluorescent intensity *I* plotted against the fluorescent dye concentration *C. R*, correlation coefficient.

### S2. Result of kymograph, plots between *a’* and *b’*, and determined velocity *v* in ELs and SMLs

The kymograph, plots between *a’* and *b’*, and determined velocity *v* in each EL and SML as obtained for the interstitial flow velocity in the whole thickness of the aorta are shown in Figs. S2.1, S2.2, and S2.3, respectively. Because the time length of the kymographs was determined as the time at which the intensity almost reached the static state, the shorter kymograph means that the concentration has early reached the steady state and that the velocity (i.e., diffusion) is high there (Fig. S2.1). The plots between *a’* and *b’* show a good and high correlation (Fig. S2.2). In Fig. S2.3, negative values are often observed in ELs and even in SMLs (0–3 from 10–13 measurements). These results indicate the low accuracy of this measurement in ELs, and therefore, the EL data was not used in further analyses. The low accuracy in ELs was attributed to the nature of the fluorescent dye; specifically, the dye was bound more easily to the ELs than to SMLs (See Supplementary Materials S4). Under such conditions, the inverse intensity gradient, where the intensity on the adventitial side is higher than that on the intimal side in the *r*-direction, can be locally seen; it results in a negative slope in the plots between *a’* and *b’*.

**Fig. S2.1.**
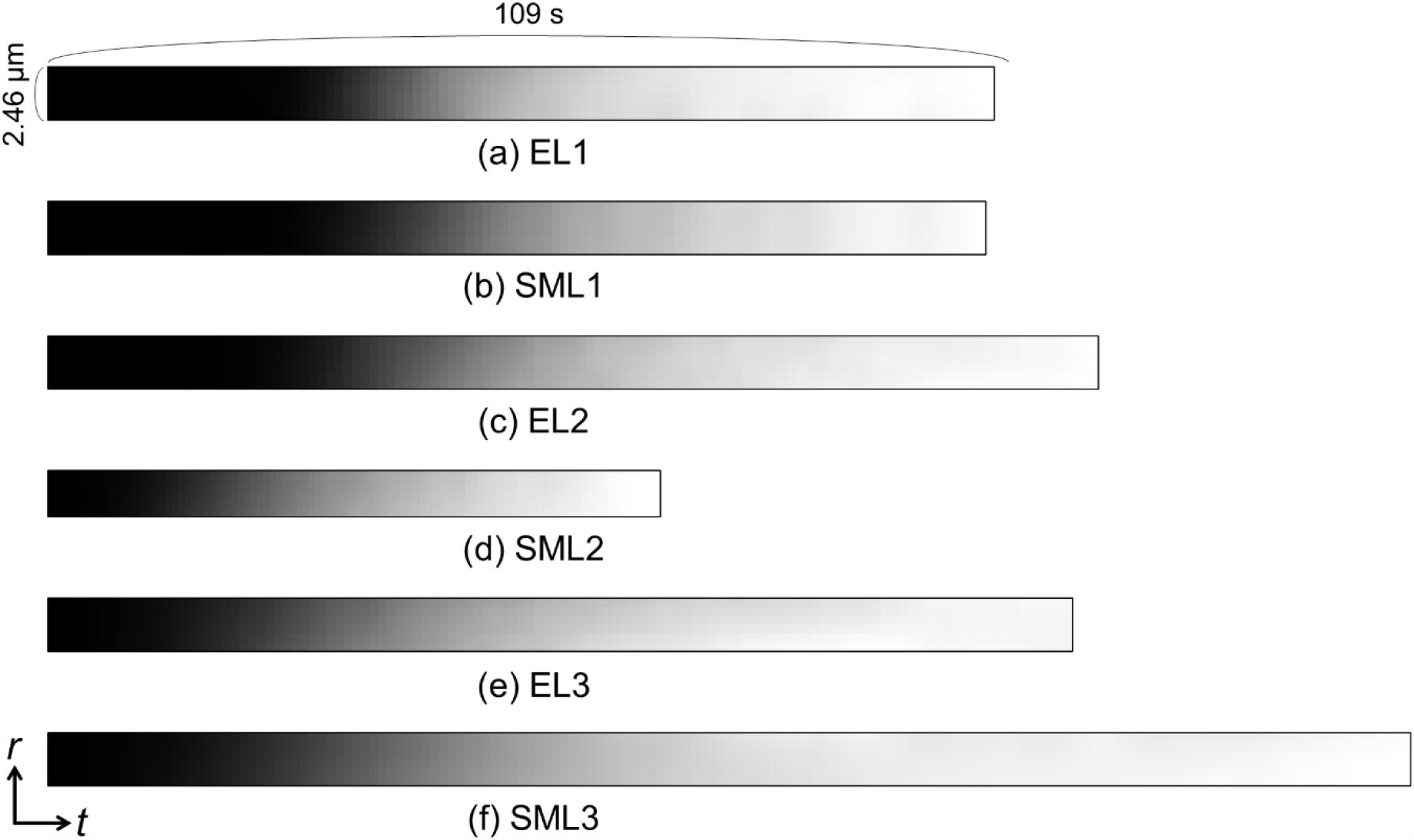
Kymograph of fluorescent dye solution in (a) EL1, (b) SML1, (c) EL2, (d) SML2, (e) EL3, and (f) SML3 from the intraluminal side at intraluminal pressure of 160 mmHg. *t*, time axis; *r*, radial axis.

**Fig. S2.2.**
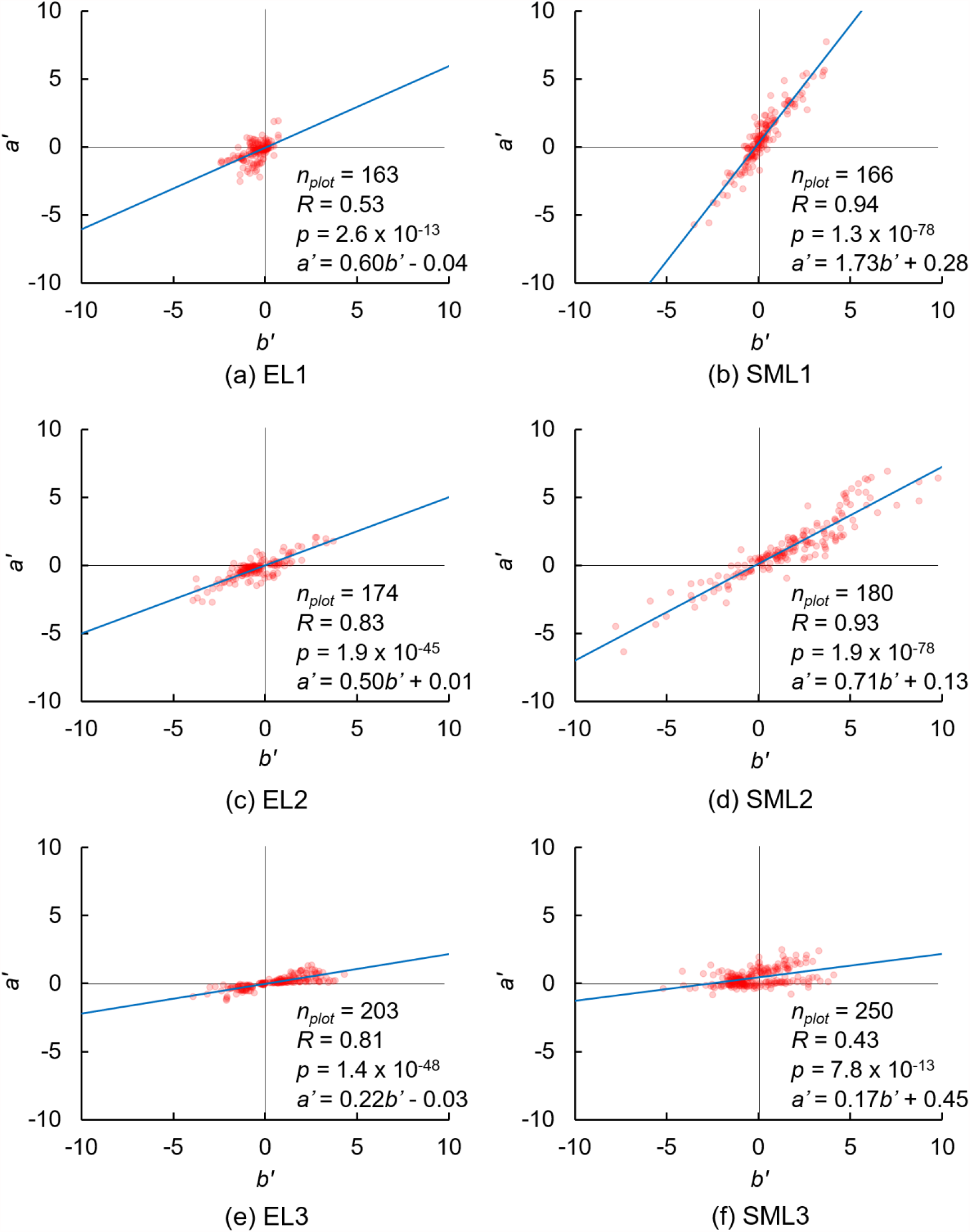
Typical plots of *a’* and *b’* in Eq. (8) in (a) EL1, (b) SML1, (c) EL2, (d) SML2, (e) EL3, and (f) SML3 from the intraluminal side at intraluminal pressure of 160 mmHg. *n*_plot_, number of plots.

**Fig. S2.3.**
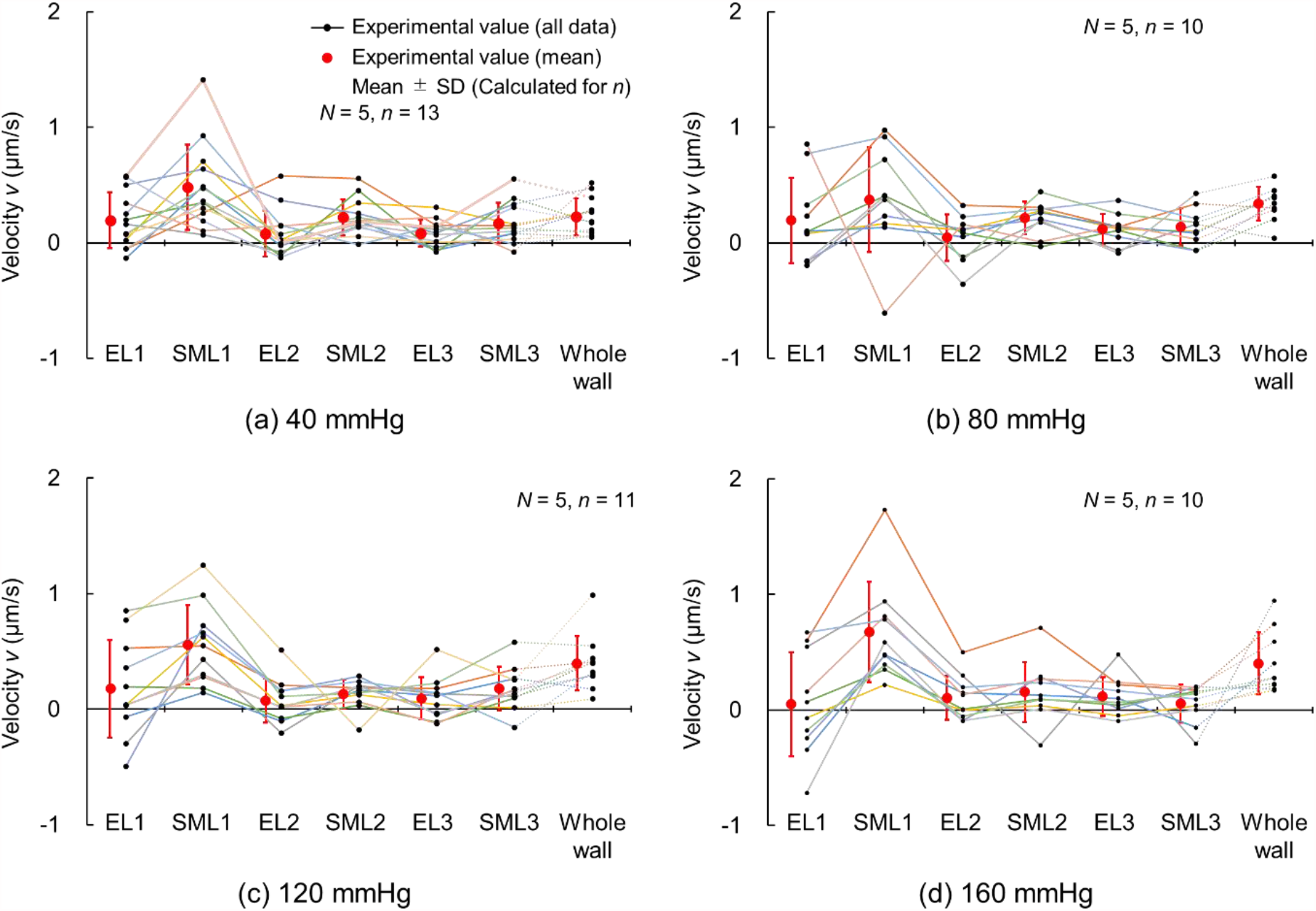
Interstitial flow velocity in local ELs and SMLs along with the velocity of the whole wall at (a) 40, (b) 80, (c) 120, and (d) 160 mmHg. *N*, Number of mice; *n*, number of data.

### S3. Changes in wall thickness during pressurization

Because the aorta is viscoelastic, the wall thickness might decrease during pressurization, resulting in an increased pressure gradient through the aortic walls and interstitial flow velocity. Here, we investigated the wall thickness at the start time *t*_*start*_ and end time *t*_*end*_ of the captured image.

The images in Fig. 3 were used for this analysis. The distance between the internal and the external ELs (*i*.*e*., *r*_*EEL*_) was determined as the wall thickness *W*. The wall thickness was measured at *t*_*start*_ and *t*_*end*_, and they were compared and tested with a paired Student’s *t*-test for all pressure levels *P*. The correlation coefficient between *W* and *P* was tested using Student’s *t*-test. The significant level was set as *p* = 0.05.

Figure S3.1 shows the wall thickness *W* at the start time *t*_*start*_ and end time *t*_*end*_. *W* at *t*_*start*_ was comparable to *W* at *t*_*end*_ for all pressure levels, and there were no significant differences between them, indicating that the wall thickness did not change during the experiment. The wall thickness significantly and negatively correlated with the intraluminal pressure (*R* = −0.507 for *t*_*start*_, *p* < 0.05; *R* = −0.505 for *t*_*end*_, *p* < 0.05) as reported in Guo et al. (Guo et al., 2007). Thus, the pressure gradient in the radial direction increased with an increase in the pressure owing to not only the pressure itself but also the reduction in the wall thickness.

**Fig. S3.1.**
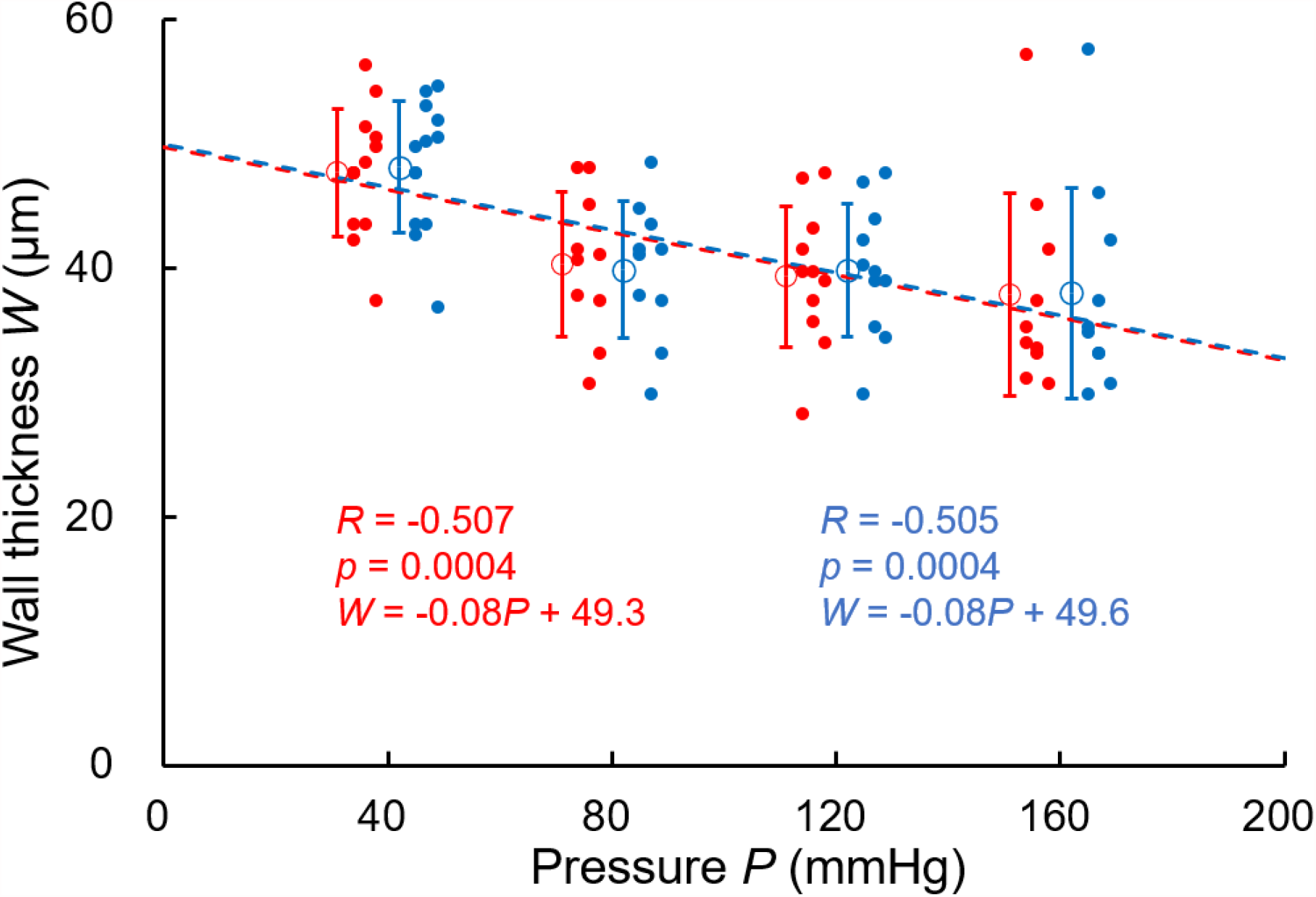
Aortic wall thickness *W* plotted to intraluminal pressure *P* at the start time *t*_start_ (red) and end time *t*_end_ (blue). Number of mice *N* = 5; number of measurement data *n* = 13 at 40 mmHg, *n* = 10 at 80 mmHg, *n* = 11 at 120, and *n* = 10 at 160 mmHg; *R*, correlation coefficient.

### S4. Why was the intensity in the EL region higher than that in the SML region?

The fluorescent intensity of the fluorescent dye solution was higher in ELs than in SMLs. This fluorescent intensity in ELs was not generated from the autofluorescence of elastin because the intensity was not high at time *t* = *t*_start_ (images in top row in Fig. 4). Thus, we hypothesized that the fluorescent dye, uranine, is more bound to ELs than to SMLs, and we evaluated this hypothesis.

Aortic samples were sliced with a microslicer as described in a previous report (Sugita et al., 2021). The samples were immersed in 1.06 mM uranine fluorescent dye solution for 5 min at room temperature and then washed with the buffer. A sample image was observed under the two-photon microscope. A 512 × 512 pixel (212 µm × 212 µm) image stack (2.00-µm intervals between image sectioning and total thickness of 40–80 µm) was captured. After taking the maximum intensity projection, the intensities in both regions were measured by tracing the regions of ELs and SMLs using ImageJ image analysis software.

Figure S4.1 shows the fluorescent images before and after immersing the specimen in the fluorescent dye solution. The intensity after immersion is clearly higher in the ELs than in SMLs. Figure S4.2 shows the quantified intensity in ELs and SMLs after immersion in the fluorescent dye solution. In this graph, the intensity is normalized with the intensity before immersion in the fluorescent dye solution to remove the effect of autofluorescence. The fluorescent intensity in ELs is significantly higher than that in SMLs. Thus, we conclude that the uranine fluorescent dye binds more to ELs than to SMLs.

**Fig. S4.1.**
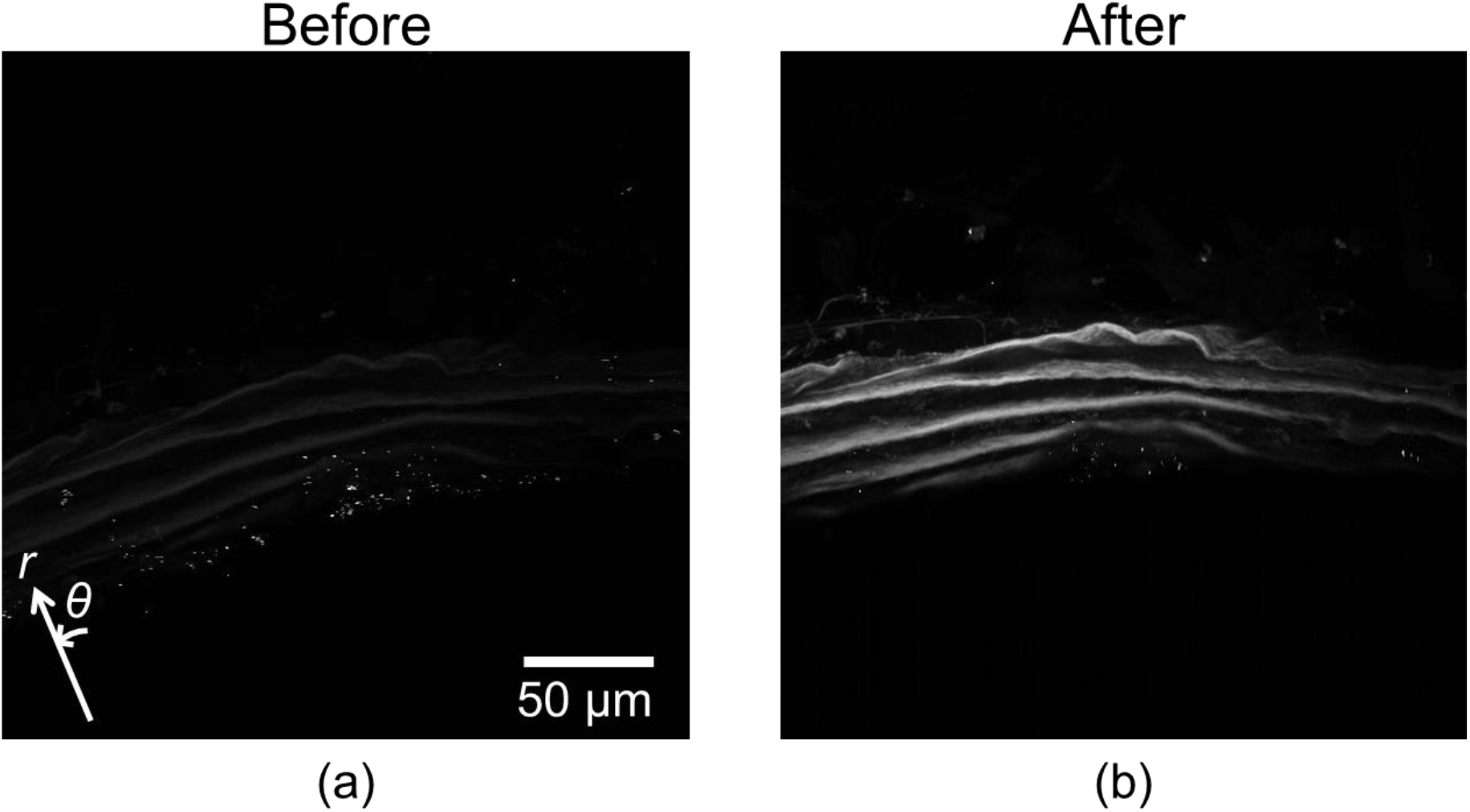
Fluorescent images of the aorta (a) before and (b) after immersion in the fluorescent dye solution. In (b), the fluorescent intensity in ELs is much higher than that in SMLs.

**Fig. S4.2.**
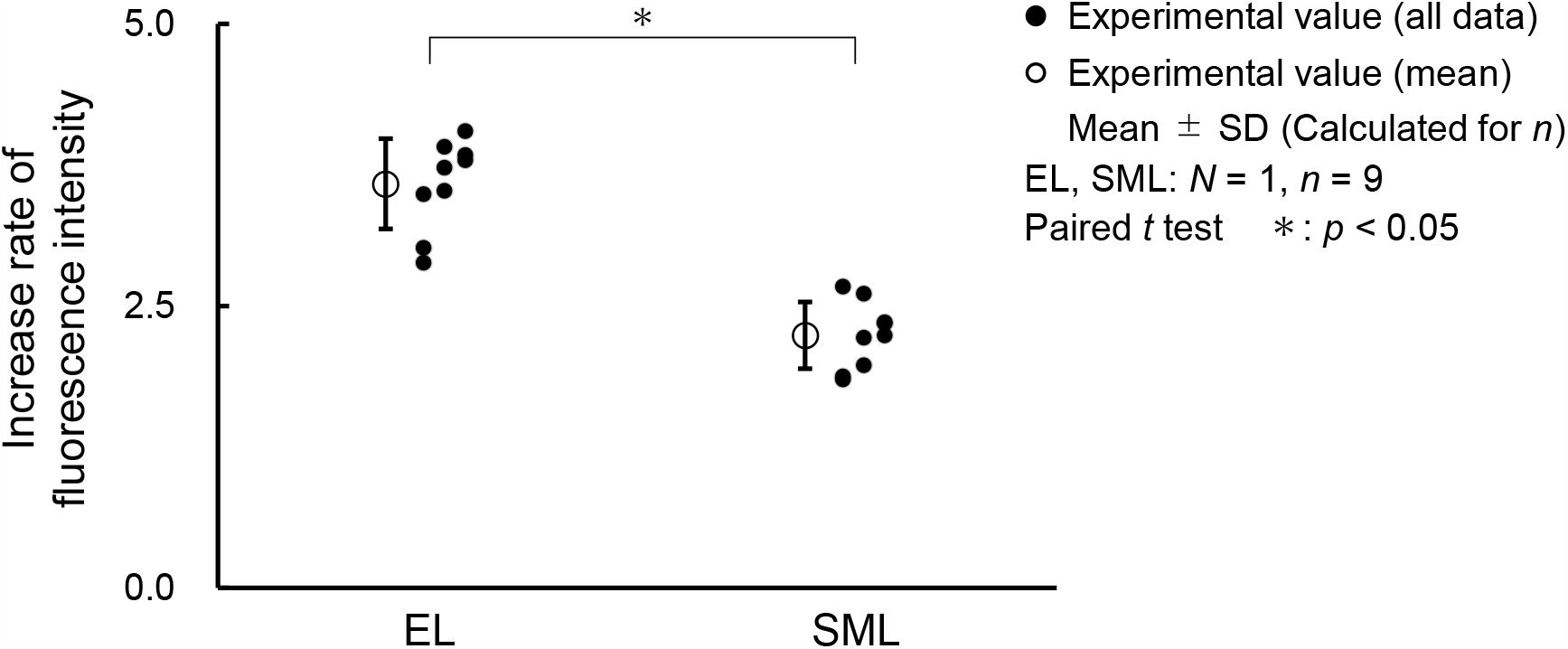
The ratio of the fluorescent intensity before immersion to that after immersion in the fluorescent dye solution in ELs and SMLs. *N*, Number of mice; *n*, number of data.

### S5. Comparison of convective velocity with diffusive velocity

Because the fluorescent dye moves by both the interstitial flow and diffusion, we compared the magnitudes of both effects. To evaluate the magnitudes of the velocity and diffusion, the Peclet number *Pe*, which shows the ratio of convective velocity to diffusive velocity, was calculated according to Baldwin et al. (Baldwin et al., 1997) as follows:

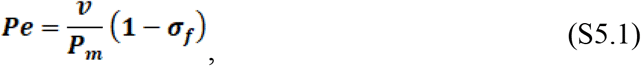

where *P*_*m*_ is the permeability coefficient and *σ*_*f*_, the filtration reflection coefficient. *Pe* > 1 indicates that the convective velocity dominates over the diffusive velocity. The diffusion coefficient *k* is obtained as the division of *P*_m_ and the wall thickness *W*:

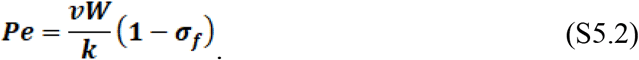

Shou et al. (Shou et al., 2006) estimated *σ*_*f*_ of trypan blue (molecular weight: 961) as 0.064. Because the molecular weight of uranine is 376, *σ*_*f*_ of uranine must be <0.064. The averaged *v* and *k* values obtained in this study are 0.33 µm/s and 0.38 µm^2^/s (40–160 mmHg), respectively, and the averaged thickness is *W* = 41.8 µm. By using *σ*_*f*_ = 0.064, *Pe* is calculated as 34. Because *σ*_*f*_ of uranine must be <0.064, the Peclet number is >34. This indicates that the diffusion velocity in the aorta is much smaller than the convective velocity.

### S6. Measurement of diffusion coefficient without flow

The proposed method can measure the diffusion coefficient as well as the flow velocity. Because the movement of the fluorescent dye by diffusion was much smaller than that by flow (see Supplementary Materials S5), we did not have confidence in the accuracy of the measured diffusion coefficient. Thus, to verify the magnitude of the diffusion coefficient, we measured the diffusion coefficient under the condition without flow.

We tried to measure only the diffusion coefficient. However, under zero intraluminal pressurization, the shape of the aorta cannot be maintained as a cylindrical pipe, resulting in a difficulty in observing the fluorescent dye. Thus, we applied an intraluminal pressure of only 5 mmHg to maintain the shape of the aortic sample and a small interstitial flow in the aorta in the experiments described in this section.

The three-way valve (no. 3 in Fig. 1) was opened, and the intraluminal pressure was kept at 80 mmHg until the fluorescent dye solution reached the aortic sample. Then, the pressure applied by the electropneumatic regulator was temporarily stopped and a hydrostatic pressure of 5 mmHg was applied to the intraluminal side of the aorta by vertically elevating the reservoir. The fluorescent dye was observed and image analysis was performed.

Figure S6.1a shows a time-lapsed image of fluorescent in the aortic wall at 5 mmHg in the radial-circumferential (*r*–*θ*) plane. The fluorescent intensity gradually increased with increasing time, indicating that the fluorescent dye moved into the aortic walls. After 200 s, the intensity in the adventitial side of the media became 0, indicating that this side was locally destroyed by laser ablation. Because the adventitial side of the media was destroyed, only the intimal side (region within 29 µm from the internal EL) was analyzed. Further, only images captured until 200 s were used because the intensity in the intimal side suddenly increased. This might have been caused by the destruction of the adventitial side, and the laser easily reached the intimal side. The average intensity in the analyzed region until 200 s (Fig. S6b) and the normalized kymograph (Fig. S6c) clearly show the increase in the fluorescent intensity with increasing time. Figure S6d shows the plots of (*a’, b’*). The fitted linear regression line was *a’* = 0.01*b’* + 0.16, and no significant correlation was observed. This result confirms that the interstitial flow was negligibly small (0.01 µm/s) at 5 mmHg. The diffusion coefficient (*k* = 0.16 µm^2^/s) almost concurred with the data at 40 mmHg (*k* = 0.24 ± 0.18 µm^2^/s). Therefore, we confirmed that the diffusion coefficients in the aortic media as measured using the proposed method were appropriate even under higher intraluminal pressures.

**Fig. S6.1.**
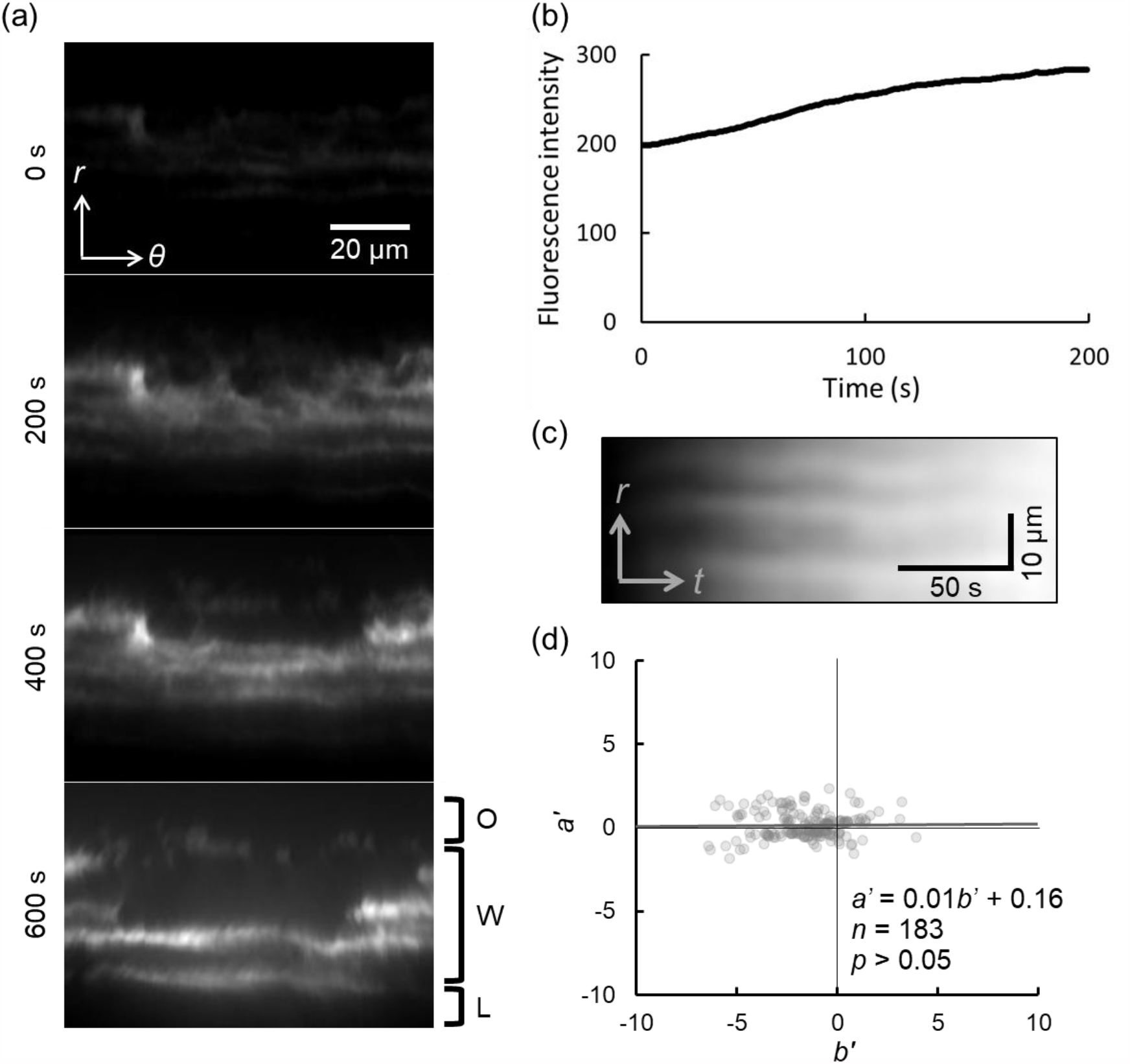
Images of interstitial flow in the aortic wall at intraluminal pressure of 5 mmHg, and image analysis processes of interstitial flow velocity and diffusion coefficient. (a) Time-lapsed images of the aortic walls in the radius-circumferential (*r*-*θ*) cross-section at intraluminal pressure of 5 mmHg. Images were captured at 0, 200, 400, and 600 s after capturing a photograph. From the top to the bottom sides, the outside of the aorta (O), aortic wall (W), and lumen (L) are shown. (b) Time-course change in the average fluorescence intensity taken on the intimal side of the aortic wall. (c) A normalized kymograph with 0 at its left edge and 1 at its right edge. The normalization was performed at each *r*-axis. *t*, time axis. (d) Plots of *a*’ and *b*’ at intraluminal pressure of 5 mmHg. *n*, number of plots.

---

**Movie 1** Time-lapsed images of ELs (red) and fluorescent dye (green) solutions in the *r*-*θ* plane at 40, 80, 120, and 160 mmHg.

## References

Baldwin, A. L., & Wilson, L. M. (1993). Endothelium increases medial hydraulic conductance of aorta, possibly by release of EDRF. Am J Physiol, 264(1 Pt 2), H26–32. doi:10.1152/ajpheart.1993.264.1.H26

Baldwin, A. L., Wilson, L. M., Gradus-Pizlo, I., Wilensky, R., & March, K. (1997). Effect of atherosclerosis on transmural convection and arterial ultrastructure. Arteriosclerosis, Thrombosis, and Vascular Biology, 17(12), 3365–3375. doi:10.1161/01.ATV.17.12.3365

Baldwin, A. L., Wilson, L. M., & Simon, B. R. (1992). Effect of pressure on aortic hydraulic conductance. Arteriosclerosis and Thrombosis: A Journal of Vascular Biology, 12(2), 163–171. doi:10.1161/01.ATV.12.2.163

Baldwin, A. L., Winlove, C. P., & Caro, C. G. (1982). Structural response of the rabbit thoracic aorta and inferior vena cava to in situ collapse. Artery, 10(6), 420–439.

Bevan, R. D. (1976). An autoradiographic and pathological study of cellular proliferation in rabbit arteries correlated with an increase in arterial pressure. Journal of Vascular Research, 13(1-2), 100–128. doi:10.1159/000158083

Chamley-Campbell, J. H., Campbell, G. R., & Ross, R. (1981). Phenotype-dependent response of cultured aortic smooth muscle to serum mitogens. J Cell Biol, 89(2), 379–383. doi:10.1083/jcb.89.2.379

Chooi, K. Y., Comerford, A., Sherwin, S. J., & Weinberg, P. D. (2016). Intimal and medial contributions to the hydraulic resistance of the arterial wall at different pressures: a combined computational and experimental study. Journal of The Royal Society Interface, 13(119), 20160234. doi:10.1098/rsif.2016.0234

Concannon, J., Dockery, P., Black, A., Sultan, S., Hynes, N., McHugh, P. E., … McGarry, J. P. (2020). Quantification of the regional bioarchitecture in the human aorta. Journal of Anatomy, 236(1), 142–155. doi:https://doi.org/10.1111/joa.13076

Dabagh, M., Jalali, P., Konttinen, Y. T., & Sarkomaa, P. (2008). Distribution of shear stress over smooth muscle cells in deformable arterial wall. Med Biol Eng Comput, 46(7), 649–657. doi:10.1007/s11517-008-0338-7

Deng, X., Marois, Y., & Guidoin, R. (1998). Fluid filtration across the arterial wall under flow conditions: is wall shear rate another factor affecting filtration rate? Ann N Y Acad Sci, 858, 105–115. doi:10.1111/j.1749-6632.1998.tb10145.x

Gaballa, M. A., Raya, T. E., Simon, B. R., & Goldman, S. (1992). Arterial mechanics in spontaneously hypertensive rats. Mechanical properties, hydraulic conductivity, and two-phase (solid/fluid) finite element models. Circ Res, 71(1), 145–158. doi:10.1161/01.res.71.1.145

Huang, Y., Rumschitzki, D., Chien, S., & Weinbaum, S. (1994). A fiber matrix model for the growth of macromolecular leakage spots in the arterial intima. Journal of Biomechanical Engineering, 116(4), 430–445. doi:10.1115/1.2895794

Kim, S. A., Sung, J. Y., Woo, C. H., & Choi, H. C. (2017). Laminar shear stress suppresses vascular smooth muscle cell proliferation through nitric oxide-AMPK pathway. Biochem Biophys Res Commun, 490(4), 1369–1374. doi:10.1016/j.bbrc.2017.07.033

Lever, M., Tarbell, J., & Caro, C. (1992). The effect of luminal flow in rabbit carotid artery on transmural fluid transport. Experimental Physiology, 77(4), 553–563. doi:10.1113/expphysiol.1992.sp003619

Liu, X., Huang, X., Chen, L., Zhang, Y., Li, M., Wang, L., … Zhang, M. (2015). Mechanical stretch promotes matrix metalloproteinase-2 and prolyl-4-hydroxylase α1 production in human aortic smooth muscle cells via Akt-p38 MAPK-JNK signaling. Int J Biochem Cell Biol, 62, 15–23. doi:10.1016/j.biocel.2015.02.009

Lu, X., Huxley, V. H., & Kassab, G. S. (2013). Endothelial barrier dysfunction in diabetic conduit arteries: a novel method to quantify filtration. American Journal of Physiology-Heart and Circulatory Physiology, 304(3), H398–H405. doi:10.1152/ajpheart.00550.2012

Matsumoto, T., & Hayashi, K. (1996). Stress and strain distribution in hypertensive and normotensive rat aorta considering residual strain. Journal of Biomechanical Engineering, 118(1), 62–73. doi:10.1115/1.2795947

Nguyen, T., Toussaint, J., Xue, Y., Raval, C., Cancel, L., Russell, S., … Rumschitzki, D. S. (2015). Aquaporin-1 facilitates pressure-driven water flow across the aortic endothelium. Am J Physiol Heart Circ Physiol, 308(9), H1051–1064. doi:10.1152/ajpheart.00499.2014

Ralevic, V., Kristek, F., Hudlická, O., & Burnstock, G. (1989). A new protocol for removal of the endothelium from the perfused rat hind-limb preparation. Circulation Research, 64(6), 1190–1196. doi:10.1161/01.RES.64.6.1190

Reyna, S. V., Ensenat, D., Johnson, F. K., Wang, H., Schafer, A. I., & Durante, W. (2004). Cyclic strain stimulates L-proline transport in vascular smooth muscle cells. Am J Hypertens, 17(8), 712–717. doi:10.1016/j.amjhyper.2004.03.673

Shi, Z.-D., Abraham, G., & Tarbell, J. M. (2010). Shear stress modulation of smooth muscle cell marker genes in 2-D and 3-D depends on mechanotransduction by heparan sulfate proteoglycans and ERK1/2. PLoS ONE, 5(8), e12196. doi:10.1371/journal.pone.0012196

Shou, Y., Jan, K. M., & Rumschitzki, D. S. (2006). Transport in rat vessel walls. I. Hydraulic conductivities of the aorta, pulmonary artery, and inferior vena cava with intact and denuded endothelia. Am J Physiol Heart Circ Physiol, 291(6), H2758–2771. doi:10.1152/ajpheart.00610.2005

Sjölund, M., Madsen, K., von der Mark, K., & Thyberg, J. (1986). Phenotype modulation in primary cultures of smooth-muscle cells from rat aorta: Synthesis of collagen and elastin. Differentiation, 32(2), 173–180. doi:10.1111/j.1432-0436.1986.tb00570.x

Stanley, A. G., Patel, H., Knight, A. L., & Williams, B. (2000). Mechanical strain-induced human vascular matrix synthesis: the role of angiotensin II. J Renin Angiotensin Aldosterone Syst, 1(1), 32–35. doi:10.3317/jraas.2000.007

Sterpetti, A. V., Cucina, A., Fragale, A., Lepidi, S., Cavallaro, A., & Santoro-D’Angelo, L. (1994). Shear stress influences the release of platelet derived growth factor and basic fibroblast growth factor by arterial smooth muscle cells. Winner of the ESVS prize for best experimental paper 1993. Eur J Vasc Surg, 8(2), 138–142. doi:10.1016/s0950-821x(05)80448-7

Sugita, S., Kato, M., Wataru, F., & Nakamura, M. (2020). Three-dimensional analysis of the thoracic aorta microscopic deformation during intraluminal pressurization. Biomechanics and Modeling in Mechanobiology, 19(1), 147–157. doi:10.1007/s10237-019-01201-w

Sugita, S., & Matsumoto, T. (2013). Quantitative measurement of the distribution and alignment of collagen fibers in unfixed aortic tissues. Journal of Biomechanics, 46(7), 1403–1407. doi:10.1016/j.jbiomech.2013.02.003

Sugita, S., & Matsumoto, T. (2017). Multiphoton microscopy observations of 3D elastin and collagen fiber microstructure changes during pressurization in aortic media. Biomechanics and Modeling in Mechanobiology, 16(3), 763–773. doi:10.1007/s10237-016-0851-9

Tada, S., & Tarbell, J. M. (2000). Interstitial flow through the internal elastic lamina affects shear stress on arterial smooth muscle cells. American Journal of Physiology-Heart and Circulatory Physiology, 278(5), H1589–H1597. doi:10.1152/ajpheart.2000.278.5.H1589

Tada, S., & Tarbell, J. M. (2002). Flow through internal elastic lamina affects shear stress on smooth muscle cells (3D simulations). Am J Physiol Heart Circ Physiol, 282(2), H576–584. doi:10.1152/ajpheart.00751.2001

Tarbell, J. M., Lever, M. J., & Caro, C. G. (1988). The effect of varying albumin concentration of the hydraulic conductivity of the rabbit common carotid artery. Microvascular Research, 35(2), 204–220. doi:10.1016/0026-2862(88)90063-5

Tedgui, A., & Lever, M. J. (1984). Filtration through damaged and undamaged rabbit thoracic aorta. Am J Physiol, 247(5 Pt 2), H784–791. doi:10.1152/ajpheart.1984.247.5.H784

Tedgui, A., & Lever, M. J. (1987). Effect of pressure and intimal damage on 131I-albumin and [14C]sucrose spaces in aorta. American Journal of Physiology-Heart and Circulatory Physiology, 253(6), H1530–H1539. doi:10.1152/ajpheart.1987.253.6.H1530

Toussaint, J., Raval, C. B., Nguyen, T., Fadaifard, H., Joshi, S., Wolberg, G., … Rumschitzki, D. S. (2017). Chronic hypertension increases aortic endothelial hydraulic conductivity by upregulating endothelial aquaporin-1 expression. American Journal of Physiology-Heart and Circulatory Physiology, 313(5), H1063–H1073. doi:10.1152/ajpheart.00651.2016

Vargas, C. B., Vargas, F. F., Pribyl, J. G., & Blackshear, P. L. (1979). Hydraulic conductivity of the endothelial and outer layers of the rabbit aorta. American Journal of Physiology-Heart and Circulatory Physiology, 236(1), H53–H60. doi:10.1152/ajpheart.1979.236.1.H53

Wada, K. (2010). Detection ofmultivariate ouliers-modified stahel-dnoho estimators-. Research memoir of the statistics, 67, 89–157.

Wang, S., & Tarbell, J. M. (2000). Effect of fluid flow on smooth muscle cells in a 3-dimensional collagen gel model. Arterioscler Thromb Vasc Biol, 20(10), 2220–2225. doi:10.1161/01.atv.20.10.2220

Wang, Z., Liu, M., Liu, X., Sun, A., Fan, Y., & Deng, X. (2019). Hydraulic conductivity and low-density lipoprotein transport of the venous graft wall in an arterial bypass. Biomedical Engineering OnLine, 18(1), 50. doi:10.1186/s12938-019-0669-7

## References

Guo, X., Lanir, Y., & Kassab, G. S. (2007). Effect of osmolarity on the zero-stress state and mechanical properties of aorta. American Journal of Physiology-Heart and Circulatory Physiology, 293(4), H2328–H2334. doi:10.1152/ajpheart.00402.2007

Sugita, S., Mizuno, N., Ujihara, Y., & Nakamura, M. (2021). Stress fibers of the aortic smooth muscle cells in tissues do not align with the principal strain direction during intraluminal pressurization. Biomechanics and Modeling in Mechanobiology. doi:10.1007/s10237-021-01427-7

